# High resolution spatio-temporal assessment of simian/human immunodeficiency virus (SHIV) evolution reveals a highly dynamic process within the host

**DOI:** 10.1101/097980

**Authors:** Alison F. Feder, Christopher Kline, Patricia Polacino, Mackenzie Cottrell, Angela D. M. Kashuba, Brandon F. Keele, Shiu-Lok Hu, Dmitri A. Petrov, Pleuni S. Pennings, Zandrea Ambrose

## Abstract

The process by which drug-resistant HIV-1 arises and spreads spatially within an infected individual is poorly understood. Studies have found variable results relating how HIV-1 in the blood differs from virus sampled in tissues, offering conflicting findings about whether HIV-1 throughout the body is homogeneously distributed. However, most of these studies sample only two compartments and few have data from multiple time points. To directly measure how drug resistance spreads within a host and to assess how spatial structure impacts its emergence, we examined serial sequences from four macaques infected with RT-SHIV_mne027_, a simian immunodeficiency virus encoding HIV-1 reverse transcriptase (RT), and treated with RT inhibitors. Both viral DNA and RNA (vDNA and vRNA) were isolated from the blood (including plasma and peripheral blood mononuclear cells), lymph nodes, gut, and vagina at a median of four time points and RT was characterized via single-genome sequencing. The resulting sequences reveal a dynamic system in which vRNA rapidly acquires drug resistance concomitantly across compartments through multiple independent mutations. Fast migration results in the same viral genotypes present across compartments, but not so fast as to equilibrate their frequencies through time. The blood and lymph nodes were found to be compartmentalized rarely, while both the blood and lymph node were more frequently different from mucosal tissues. There is some evidence for an increase in compartmentalization after the onset of selective pressure. This study suggests that even oft-sampled blood does not fully capture the viral dynamics in other parts of the body, especially the gut where vRNA turnover was faster than the plasma and vDNA retained fewer wild-type viruses than other sampled compartments. Our findings of transient compartmentalization across multiple tissues may help explain the varied results of previous compartmentalization studies in HIV-1.

**Author Summary:** HIV-1 is difficult to treat because the virus can evolve to become drug-resistant within the body, but we have an incomplete understanding of where drug resistant viruses originate and how they spread within a person. In this study, four macaques were infected with RT-SHIV, a simian immunodeficiency virus with an HIV-1 reverse transcriptase coding region, which can be targeted with standard HIV drugs. We sampled virus from the macaques before, during and after they became resistant to administered drugs and determined the genetic viral sequences in several parts of the body: blood, lymph nodes, gut, and vagina. We found that drug resistance emerged across compartments nearly simultaneously, and drug resistance evolved multiple independent times within each macaque. Although migration of RT-SHIV between compartments is fast, compartments do not have the same distribution of viral genotypes. This is important because although studies typically sample virus from the blood to study how HIV-1 evolution in humans, our study suggests that it is not fully representative of other parts of the body, particularly the gut.

## 1. Introduction

The development of drug resistance in human immunodeficiency virus type 1 (HIV-1) to antiretroviral therapies represents a major public health obstacle. The prevalence of drug resistance in many regions of the world is now above 5% and increasing, driven in part by drug resistance transmitted from one infected individual to another (1). Transmitted drug resistance has been observed at frequencies between 2.8% and 11.5% in different regions of the world, rendering standard first line therapies ineffective and necessitating costlier therapies for treatment-naïve patients. Despite the increasing prevalence of drug resistance, we still have an incomplete understanding of how drug resistance emerges and spreads spatially within a patient.

A comprehensive understanding of the drug resistance spatial landscape, such as when and where drug resistance arises and how it spreads within a patient, could inform administration practices of current therapy and the design of new antiretroviral therapies. Previous studies have suggested that HIV-1 drug resistance originates primarily in tissues with low or absent drug concentration (i.e., drug sanctuaries) (2-4). If this holds true, then efforts should focus on improving drug penetration in these tissues.

Perhaps more importantly, understanding how drug resistance spreads within the body could improve understanding of the ecology of the virus within a patient, which, in turn, could help determine the best way to sample the viral population, monitor transmission chains, and design eradication strategies. Although nearly all sampling of HIV-1 is done in the blood plasma, it is unclear how well the plasma represents the broader HIV-1 population. Differences between viruses from plasma and tissues could drive misdiagnoses of pre-existing drug-resistant variants, resulting in incorrectly applied therapies and selection for further drug resistance. If virus in the genital tract forms a different population than virus circulating in the blood, this should be taken into account when sampling in order to allow better reconstruction of transmission chains and epidemiological tracking of transmitted drug resistance. Finally, understanding HIV-1 genetic variation across the body might contribute to our understanding of viral reservoirs and the potential for virus eradication.

It has long been recognized that it is important to understand the relationship between virus in the blood and different tissues and many studies have examined this. However, the results are equivocal and some studies find evidence for compartmentalization, whereas others do not (see Table 1 for an overview). Nearly all studies that compared compartments across time found that compartments were only different transiently (5-9). The majority of studies with multiple time points examined the relationship among different T cell subsets in the blood (7, 9, 10), and it is unclear whether these patterns hold in other comparisons, including tissues. Most studies that have been conducted focused on the comparison between blood and only one type of tissue though (see (11-13)), and thus cannot give us information on the relationships between several compartments. Taken together, previous studies suggest the presence of a complicated spatial landscape on which viral evolution occurs, and much still needs to be done to connect not only how other compartments relate to the blood plasma, but also how they relate to each other.

**Table 1:** Summary of results from different HIV-1 compartmentalization studies.

|  | **Evidence mostly against compartmentalization** | **Evidence both for and against compartmentalization** | **Evidence mostly for compartmentalization** |
| --- | --- | --- | --- |
| Plasma v. Lymph Node | Boritz et al., 2016 (56)<br>Josefsson et al., 2013 (57) |  |  |
| Plasma v. Female Genital Tract |  | Bull, et al., 2009 (58)<br>Bull, et al., 2013 (5)<br>Kelley, et al., 2010 (6) | Philpott, et al. 2005 (59) |
| Plasma v. Male Genital Tract/Semen |  | Diem, et al., 2008 (60) |  |
| Plasma v. Central Nervous System |  | Schnell, et al., 2009 (61)<br>Schnell, et al., 2010 (8)<br>Sturdevant, et al., 2012 (62)<br>Ritola, et al., 2005 (63) | Smit, et al., 2004 (64)<br>Wong, et al., 1997 (65)<br>Ohagen, et al., 2003 (66) |
| Plasma v. Gut | Imamichi, et al., 2011 (11) | Zhang, et al., 2002 (67)<br>Lerner, et al., 2011 (68)<br>Lewis, et al., 2013 (69)<br>Van Marle, et al., 2010 (70)<br>Rozera, et al, 2014 (71) | van Marle, et al., 2007 (72) |
| Among different T cell populations | Heeregrave et al, 2003 (10) | Boritz et al., 2016 (56)<br>Josefsson et al., 2013 (57)<br>Delobel, et al., 2005 (73)<br>Potter, et al., 2004 (74)<br>Fulcher, et al., 2004 (7) | Potter, et al., 2006 (9) |

To increase our understanding of how drug resistance spreads through space and how different compartments in a host are connected to each other, we present comprehensive single-genome sequence (SGS) data from four different compartments and the blood, at four (median) time points, in four macaques infected with a simian immunodeficiency virus containing HIV-1 reverse transcriptase (RT-SHIV) (14, 15). The macaques were treated with monotherapy to induce the emergence of drug resistance, and we sampled viral RNA (vRNA) from plasma and four different compartments (lymph node, peripheral blood mononuclear cells, vagina, and gut) before, during, and after the detection of drug-resistant variants in the blood. Like HIV-1, RT-SHIV is a lentivirus with a DNA intermediate stage after infection of host cells, and we also sampled reverse transcribed viral DNA (vDNA) from the same four compartments.

These sequence data give an unprecedented look at how HIV-1 drug resistance spreads within a host. We find that drug-resistant viruses spread simultaneously across many compartments. Furthermore, drug resistance arises independently multiple times within each macaque (evidence of soft sweeps). Viruses harboring the same genotypes are often found in all compartments, but they are not always found at the same frequencies in different compartments. This suggests that although migration within these macaques is fast, it is not fast enough to create populations that are entirely well-mixed. In addition, our data suggest that the viral response to selective pressures (here, antiretroviral drugs) may transiently lead to bigger differences in population composition between compartments. Our data allowed us to observe that the viral population composition from the blood and lymph nodes appear to be more similar than among mucosal tissue samples or between blood and mucosal tissues. Samples from blood may therefore not be wholly representative of viruses in other compartments, particularly the gut, which had a faster rate of vRNA turnover, and retained less wild-type vDNA than other compartments. We also use our data to estimate two important population genetic parameters: the effective population size (N_e_), and the selection benefit conferred by drug resistance (s). These estimates will help facilitate the application of evolutionary theory to understanding how, where, and when HIV-1 drug resistance emerges and spreads.

## 2. Results

### 2.1 Experimental Design

Four macaques were infected with RT-SHIV. To intentionally select drug-resistant virus, macaques were treated with daily monotherapy beginning at 12 weeks post-infection (approximately 84 generations). Two macaques were treated with the nucleoside analog (NRTI) emtricitabine (FTC; macaques T98133 and A99039) and two received the non-nucleoside RT inhibitor (NNRTI) efavirenz (EFV; macaques A99165 and A01198). Treatment was administered for 8 weeks, followed by a 6 week treatment interruption, and then combination therapy: FTC, tenofovir (TFV), and EFV (macaques A99165 and A01198) or tenofovir, L870812, and DRV/r (macaques T98133 and A99039). RT-SHIV populations were sampled three times during monotherapy for each macaque (weeks 13, 15/16 and 20), once after treatment interruption (week 26), and once after 12-18 weeks of combination therapy (between weeks 38-44).

Samples were taken at the different time points from different blood, LN, gut, and vagina). Reverse transcriptase (RT) was sequenced from vRNA and vDNA using SGS, with a median of 28 sequences per sample (among samples with sequences) and 770 nucleotides per sequence (Supplemental Table S1). We did not find that diversity changed significantly over time except in the vRNA of A01198 and the vDNA of A99165 (Figure S1, Table S2), and values of average pairwise diversity (π) ranged between 0.2 and 0.6%, similar to what is seen in HIV-1 at a comparable amount of time after infection (16).

### 2.2 Drug Resistance Emergence and Persistence during and after Monotherapy

#### 2.2.1 Overview of drug resistance emergence in plasma viral RNA over time

In all animals, after challenge with RT-SHIV, but prior to the start of monotherapy, plasma viremia increased and a set point was reached at approximately 4 weeks post-infection (10^5^-10^6^ viral copies/mL plasma, Figure 1). After initiation of monotherapy, plasma viremia transiently decreased in all animals (decrease ranging from 33- to 566-fold) before rebounding to near pre-therapy levels. In both FTC-treated macaques (T98133 and A99039), the expected drug-resistant mutations M184V/I emerged between weeks 13-15 (1-3 weeks post-treatment), concomitantly with viral rebound and FTC concentrations in the plasma and tissues (Figure S2A). The rate at which viruses carrying DRMs displaced wild-type (WT) viruses suggests a selection coefficient for M184V/I of 0.74 in T98133 and 0.41 in A99039 (i.e., relative fitness benefits of 74% and 41%, respectively). In the EFV treated macaques however, the expected drug-resistant mutation (K103N) emerged but did not reach fixation in one macaque (A99165) and did not arise at all in the other macaque (A01198). In the animal in which K103N arose (A99165), plasma viremia initially declined but rebounded around week 15, coinciding with the spread of resistance. We suspect that the expected resistance mutation did not fix in A99165 and A01198 because selection pressure was weak due to the rapid metabolism of NNRTIs in macaques (Figure S2B), which we have observed previously (17, 18). Because EFV was not present at consistently high concentrations, we did not estimate s for K103N.

**Figure 1:**
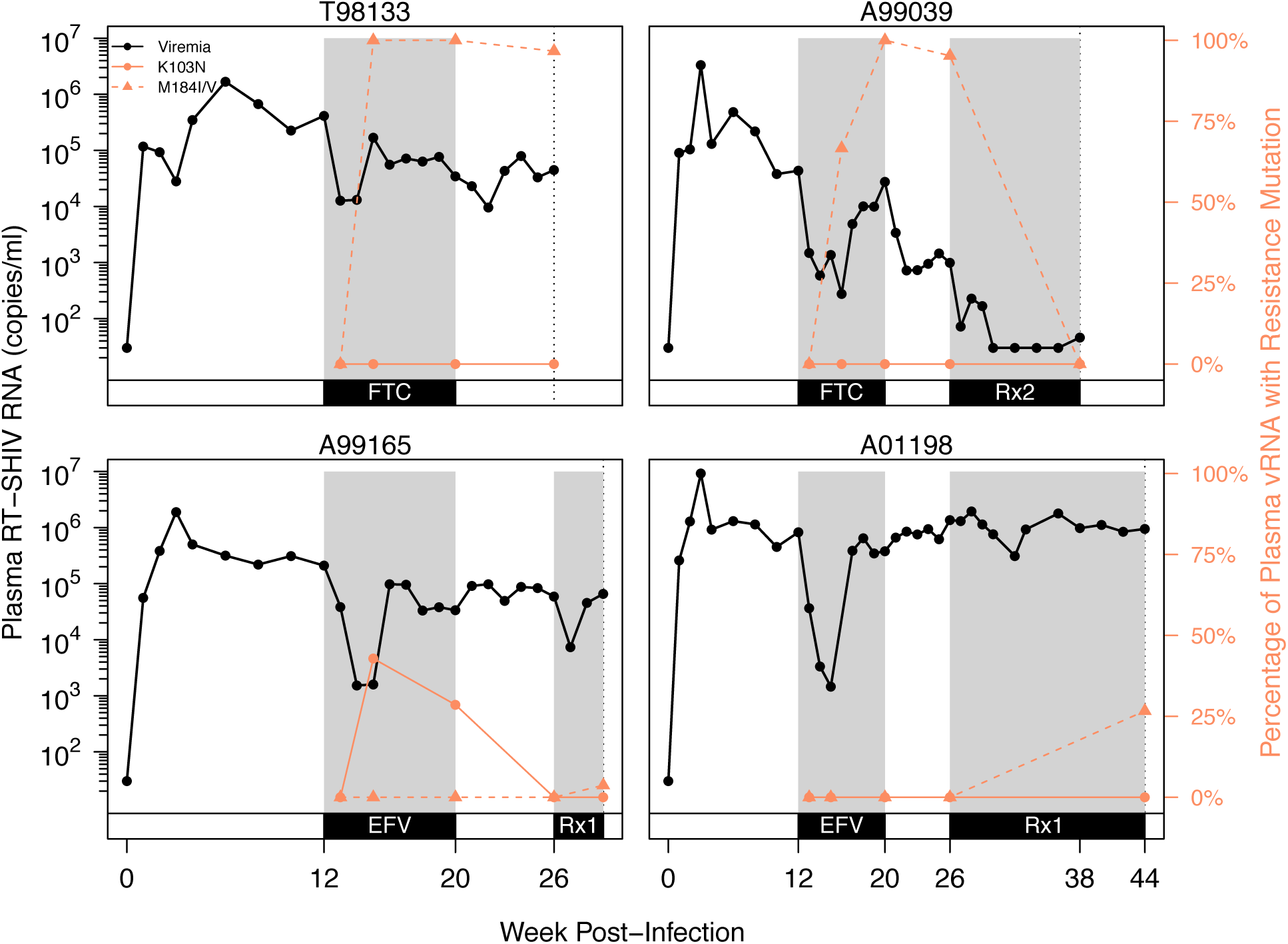
Plasma viremia and drug resistance in four macaques over the course of infection. Plasma RT-SHIV (black) and frequency of plasma vRNA with resistance mutations (orange) are plotted over time for each animal, as determined by SGS. Orange lines show the frequency of K103N (solid) or M184I/V (dashed) in the plasma vRNA. Gray shading denotes administration of drug(s) between weeks 12-20 and weeks 26-44. Black dotted line indicates the final time point for each animal. Rx1 is treatment FTC+TFV+EFV and Rx2 is the treatment TFV+L870812+DRV/r.

After monotherapy treatment was interrupted, drug-resistant plasma vRNA persisted between weeks 20-26 in both FTC-treated macaques (T98133 and A99039) (Figure 1), which suggests that the cost of drug resistance in the absence of the drug is insufficient for it to notably decrease in frequency over 6 weeks. In the EFV-treated macaque A99165, drug resistance was not observed in the plasma at the end of the treatment interruption, but in this case, the decline in frequency had already started at week 20, likely due to a decrease in EFV concentrations in plasma and tissues. Drug resistance remained absent in A01198.

Combination therapy suppressed RT-SHIV in A99039, and no further drug resistance emerged. In this macaque the M184V/I drug resistance mutations were undetectable in plasma samples taken during virologic suppression (week 38), which suggests that the cost of FTC resistance in the absence of drug is significant on this timescale or that the viruses that were sampled when VL was suppressed stem from a reservoir that had always remained WT. A99165 and A01198 both began combination therapy that included FTC, but VL was not suppressed in either animal. Drug resistance to FTC emerged (in the form of M184I and M184V, respectively), but did not fix in either macaque, reaching frequencies of 4% and 27% in the plasma, respectively.

#### 2.2.2 Drug Resistance Emerges Simultaneously in Viral RNA from Multiple Tissues

In a clinical setting, drug resistance is assessed using samples taken from the blood plasma, but how representative these samples are to levels of drug resistance in vRNA in PBMC and tissues is unclear. Here, we compared the kinetics and composition of drug-resistant variants across multiple anatomic compartments (blood, lymph node, gut, and vagina). Across the three monkeys in which drug resistance was observed, drug resistance mutations (DRMs) occurred at similar time points in the plasma and other compartments (Figure 2). In T98133, all compartments had drug-resistant vRNA detected at approximately the same time point (week 15/16). A99039 had drug-resistant vRNA first in the plasma at week 15, and then only later in the gut (week 20) and the lymph node (week 27). Although we have no vRNA samples from the vagina in this animal, the vDNA from the vagina contained drug resistance at week 15, suggesting that M184I/V emerged early in that compartment. In A99165, drug resistance was observed first at low frequency in the lymph node at week 13 and then simultaneously in gut and plasma vRNA two weeks later. We also observed that a greater proportion of vDNA from the gut contained DRMs than vDNA from other compartments (Figure 2).

**Figure 2:**
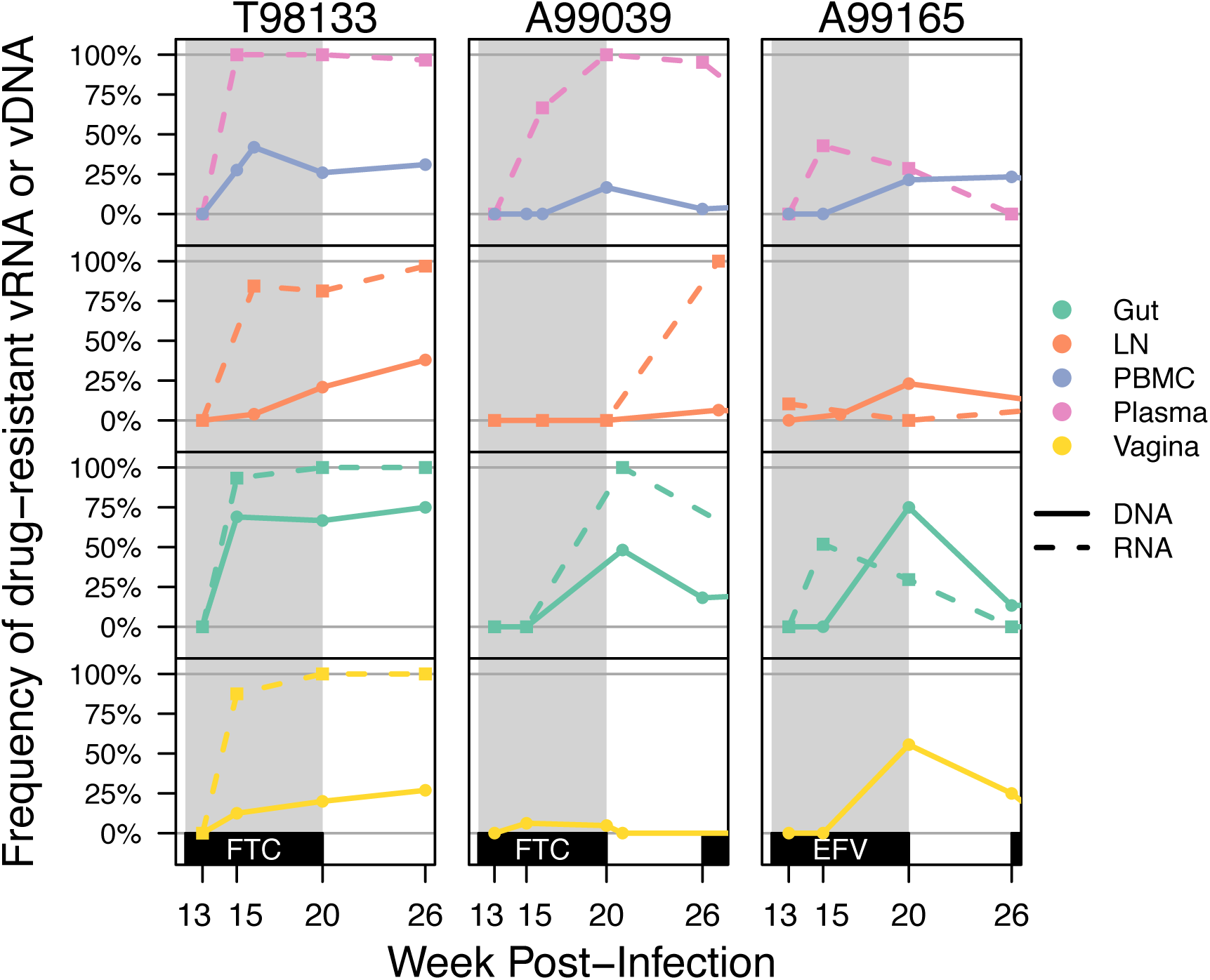
Drug resistance increase over time across all compartments. Drug resistance was defined as the proportion of vRNA (dashed lines) or vDNA (solid lines) sequences having K103N and/or M184V/I within each compartment for animals T98133, A99039, and A99165. Gray shading indicates the FTC or EFV treatment periods for the macaques.

Not only did drug resistance emerge at similar times across different compartments, but identical viral genotypes were observed at similar times in different compartments. Figures 3, 4 and S3 show the frequencies of different viral genotypes throughout time in different compartments in each of the three monkeys that had drug resistance (T98133, A99165, and A99039, respectively). Each color represents a different genotype and the WT is depicted in grey. Genotypes with drug resistance mutations are hatched. This visualization clearly shows that genotypes are usually not restricted to single compartments, but they are observed in several compartments.

**Figure 3:**
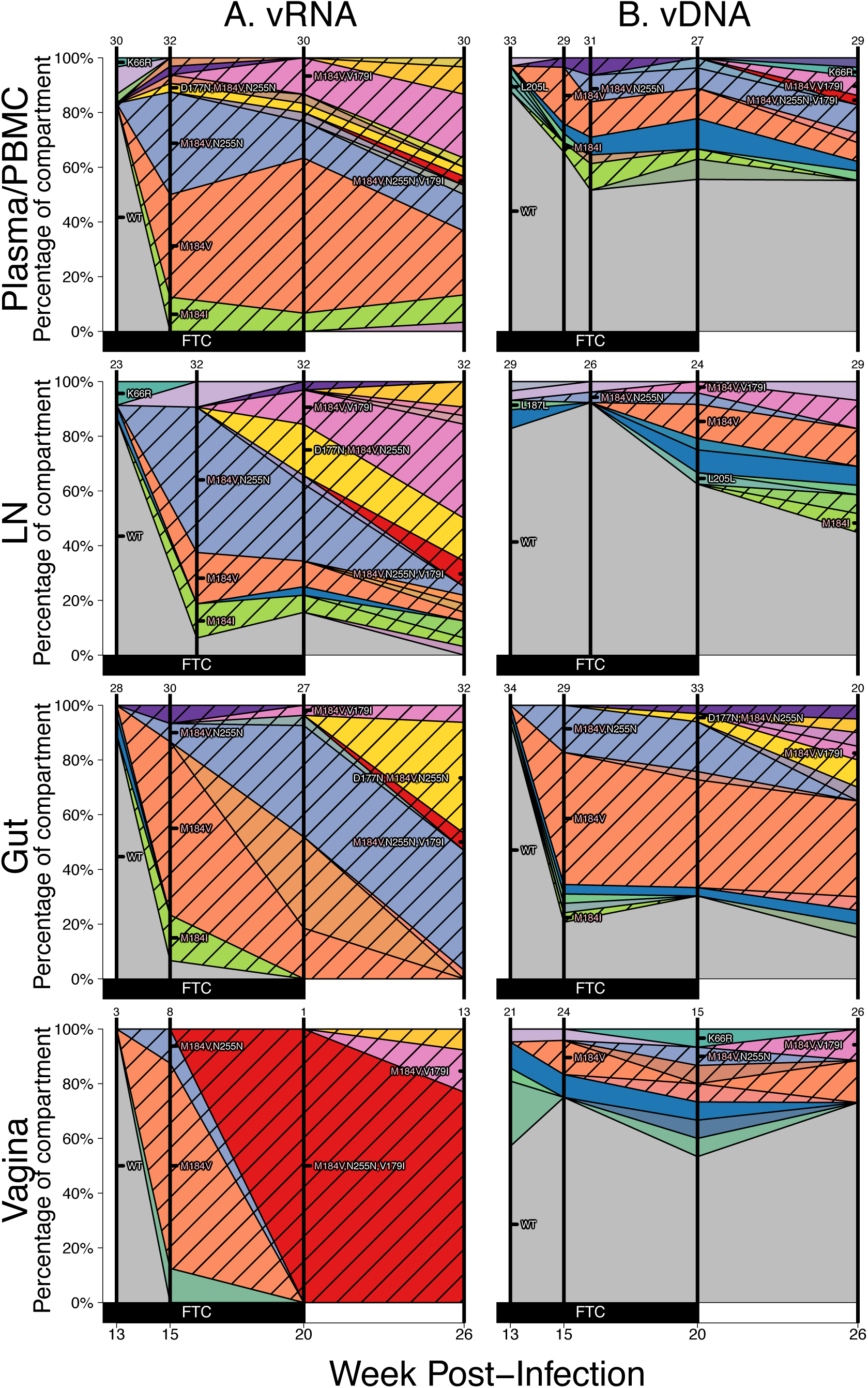
Drug resistance spreads dynamically across several compartments in macaque T98133 over the course of treatment. Colored polygons show the prevalence of different genotypes of RT-SHIV RNA (A) or DNA (B) over time in different compartments. Vertical distance at black vertical lines indicates the frequency of the sample with an exact set of mutations (i.e., genotype) at the time of sampling. Grey indicates wild-type virus, and striations on the background of a color indicate a drug-resistant genotype. Common mutations are labeled, and known drug resistance mutations are labeled in pink. For plasma/PBMC, plasma vRNA and PBMC vDNA are shown. Two sequences are considered the same genotype when all mutations found above frequency 1% in the macaque are shared. Sample sizes for at each sampling point are given above each vertical line.

**Figure 4:**
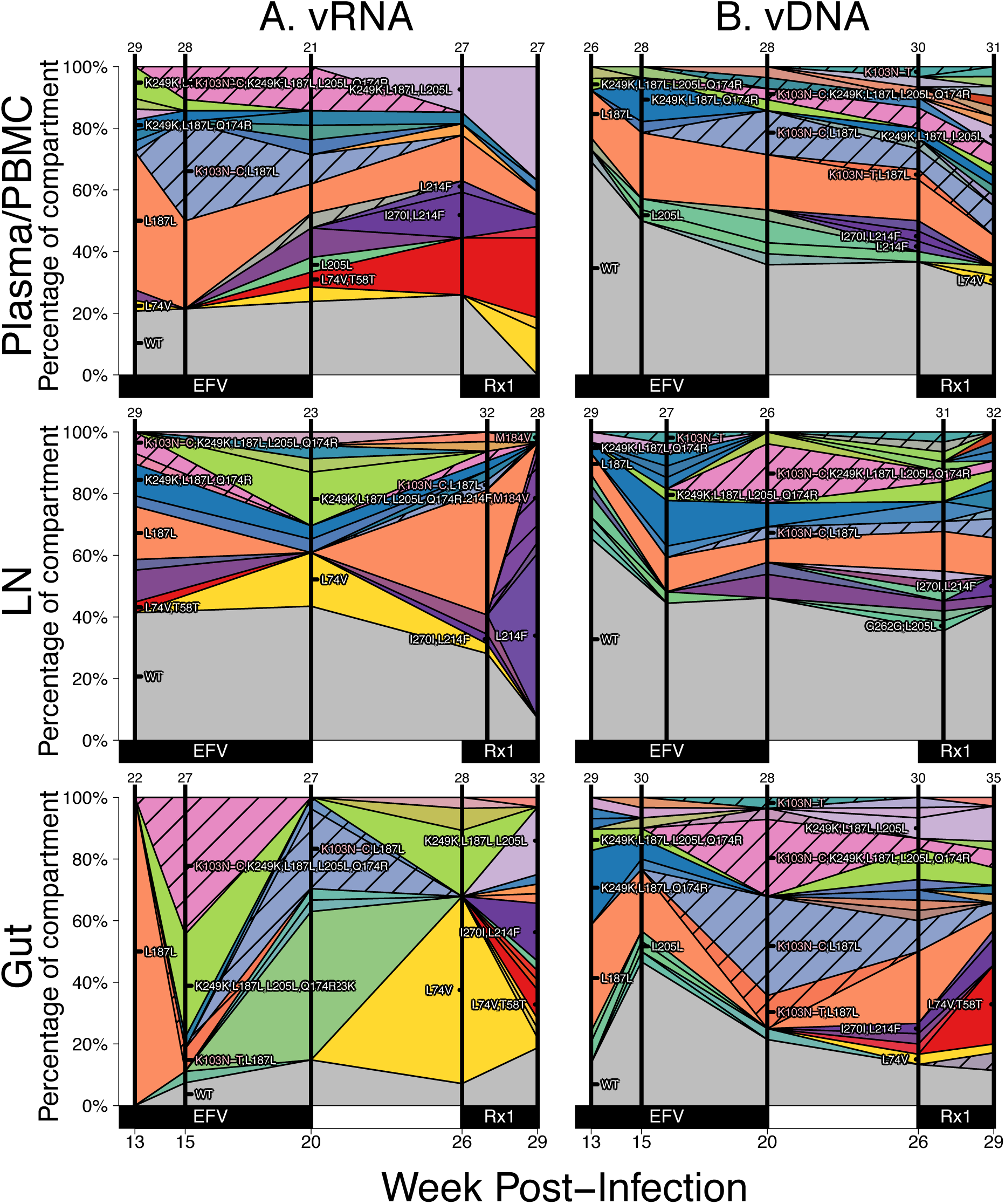
Drug resistance spreads dynamically across several compartments in macaque A99165 over the course of treatment. Figure caption the same as Figure 3. Note, except for WT, the coloration is not preserved between figures (i.e., the yellow in Figure 3 does not represent the same genotype as the yellow in Figure 4). Rx1 is treatment FTC+TFV+EFV.

For each monkey, the most frequent drug-resistant genotypes are often observed across vRNA from all compartments. For example, in T98133, at week 15/16 after initial infection, vRNA sampled from the plasma, lymph node, gut, and vagina all had two different genotypes with DRM M184V (one genotype with only M184V in orange and one genotype with an additional synonymous substitution at the 255^th^ amino acid of RT, N255N, in light blue) (Figure 3A). Three of the four compartments also contained vRNA with DRM M184I. The similarity of vRNA sequences found in each compartment, particularly vRNA with synonymous mutations (i.e., M184V linked to N255N), suggests that migration was responsible for spreading viral genotypes from one compartment to another, as opposed to independent mutational origins of each genotype in each compartment. The frequencies of genotypes remain similar between the compartments over time (Figure 3A), suggesting that substantial migration was ongoing. Similarly, in A99039 and A99165 (Figures 4A and S3A, respectively), the same drug-resistant genotypes (including linked mutations) are observed at similar time points in the vRNA in different tissue compartments.

#### 2.2.3 Drug-Resistance Mutations (DRMs) Develop on Multiple Viral Genomes

If mutation generates DRMs fast enough (mutation rate multiplied by population size larger than 1), we would expect multiple independent origins of DRMs on multiple backgrounds that spread simultaneously in what is known as a soft sweep. Alternatively, if DRMs arise only rarely, a single DRM may increase to 100% frequency in the population due to selection before a second DRM arises on a different background in the population. This is called a hard sweep (19).

There are two methods we can use to determine how many times drug resistance developed on an individual viral genome in the macaques. First, we can determine whether different DRMs (distinguishable at either the amino acid or nucleotide level) are observed simultaneously in the same population. Second, by SGS we can determine whether the same DRM occurred on different genetic backgrounds. If drug resistance increases in frequency due to a hard sweep, we should see only one DRM on one genetic background spreading (although the mutational background can later diversify due to development of new mutations). If drug resistance increases in frequency due to a soft sweep, we may see several different DRMs or we may see only one DRM that is present on different genetic backgrounds.

We first observe that drug resistance occurs multiple times through mutations distinguishable from their encodings (i.e., the DRMs themselves have different sequences). In FTC-treated macaques, RT-SHIV developed both M184V and M184I, similar to what has been observed in HIV-infected individuals (20, 21). In both cases, M184V was more common than M184I: 91.5% and 97.0% of drug-resistant vRNA variants had M184V in T98133 and A99039, respectively (Figure 3 and S3), which is similar to what has been shown in HIV-infected individuals due to a fitness advantage of M184V over M184I (21). The macaque treated with EFV (A99165) only acquired one drug resistance mutation (K103N). However, K103N can be caused by mutations either from A to U or from A to C at the third residue of amino acid 103 and can both be present in HIV-infected individuals (22). Both of these mutations were observed within A99165 (Figure 4). In both of the above cases, the encoding of two different amino acid mutations (M184V/I) or the two different nucleotide-level encodings of the same amino acid (K103N) demonstrates that at least two independent mutations must have occurred within each of the three drug-resistant macaques. This proves that soft sweeps are occurring and shows that the mutation rate multiplied by the effective population size is larger than 1 in all three monkeys.

However, SGS allowed further resolution to determine the number of times each mutation arose. In T98133, DRM M184V was seen at high frequency both with and without a synonymous N255N mutation (Figure 3). This suggests a second origin of drug resistance in which the M184V mutation occurred on a background that already had the N255N mutation. In principle, the N255N mutation may have arisen shortly after drug resistance began to spread, but given that the M184V+N255N lineage reaches a high frequency, it is more likely that M184V occurred on a genotype that already had the N255N mutation. Combined with the observation of M184I, at least three origins of resistance were likely in T98133.

In A99039, the DRM M184V occurred on three distinct backgrounds (Figure S3): 1) with L187L+M245T, 2) with L205L+R277K, and 3) with K223K+L109L+V75L. As M184I was also observed at low frequency, at least four origins of resistance were likely in A99039.

In A99165, the DRM K103N emerged on three distinguishable backgrounds (Figure 4): 1) with no other high frequency mutations, 2) with a synonymous L187L mutation, and 3) with four non-DRMs (K249K, L187L, L205L, Q174R). All three of those backgrounds were observed before detection of K103N. Further, both the nucleotide level encodings of K103N (A→U and A→C) were observed on the L187L background, suggesting at least four origins of resistance were likely in A99165.

We used the occurrence of multiple DRMs on different backgrounds to estimate the effective population size (following (23)). The idea behind this approach is that the number of times a DRM occurs in a population depends on the size of the population. In a large population, we expect DRMs to occur more often than in smaller populations. We differentiated independent origins by examining drug-resistant viral sequences with different codons at a particular DRM (e.g., K103N encoded by AAU or AAC) and different identities by haplotype (e.g., a DRM linked to another high frequency mutation), and calling the resulting estimates N_e_,_DRM_ and N_e_,_hap_, respectively. We performed both these estimates for just vRNA and combined vRNA and vDNA. We find estimates of N_e_ among the vRNA between 1.17*10^4^ and 5.71*10^5^ depending on the macaque and which technique was used (Table 2). These estimates were between 2 and 3 orders of magnitude higher than estimates using standard diversity based methods (N_e,d_).

**Table 2:** Estimates of *N_e_* taken after the spread of RT-SHIV drug resistance using three different methods.

| | $N_{e,d}$<br>**(Plasma)** | $N_{e,DRM}$<br>**(DR vRNA)** | $N_{e,hap}$<br>**(DR vRNA)** | $N_{e,DRM}$<br>**(DR all)** | $N_{e,hap}$<br>**(DR all)** |
| --- | --- | --- | --- | --- | --- |
| T98133 | $8.51 \cdot 10^1$ | $1.17 \cdot 10^4$ | $5.31 \cdot 10^4$ | $1.08 \cdot 10^4$ | $4.72 \cdot 10^4$ |
| A99165 | $7.46 \cdot 10^2$ | $4.14 \cdot 10^4$ | $2.38 \cdot 10^5$ | $6.15 \cdot 10^4$ | $3.29 \cdot 10^5$ |
| A99039 | $1.46 \cdot 10^3$ | $1.47 \cdot 10^4$ | $5.71 \cdot 10^5$ | $9.61 \cdot 10^3$ | $5.62 \cdot 10^5$ |
| A01198* | $1.73 \cdot 10^2$ | ND | ND | ND | ND |

**ND: Not done**
**\*Because no drug resistance arose in A01198, the number of origins of resistance could not be used to estimate effective population size.**

### 2.3 Frequencies of Common Viral Variants in Viral RNA suggest compartments are transiently non-well mixed

We have shown that the same genotypes containing DRMs appeared simultaneously across multiple compartments, suggesting a substantial amount of migration. However, these same genotypes are found at different frequencies in different compartments (e.g., the frequency of M184V+N255N across compartments in Figure 3). We asked whether the observed differences are due to real differences between the compartments or whether they are due to random sampling effects and fairly small sample sizes. In this context, we define two compartments to be **compartmentalized** if their vRNA genotype frequencies are significantly different when using a randomization test, and **well-mixed** otherwise.

To assess the extent of compartmentalization, we compared the frequency of vRNA genotypes in different compartments using a variance partitioning statistic, *K_ST_*. We computed *K_ST_* between all pairs of compartments sampled at the same time point (±1 week) in each macaque. Compartments needed at least 3 vRNA sequences to be included. The significance of the *K_ST_* test was assessed with a permutation test and multiple testing corrections. In 48.9% (45/92) of pairwise tests, we reject that the compartments are well-mixed (Figure 5). With a 5% false discovery rate, we expect approximately 2.5 of those tests to be false positives. This result shows that, even though DRMs appear to spread quickly from one compartment to another, migration is not pervasive enough to make the populations entirely well-mixed.

**Figure 5:**
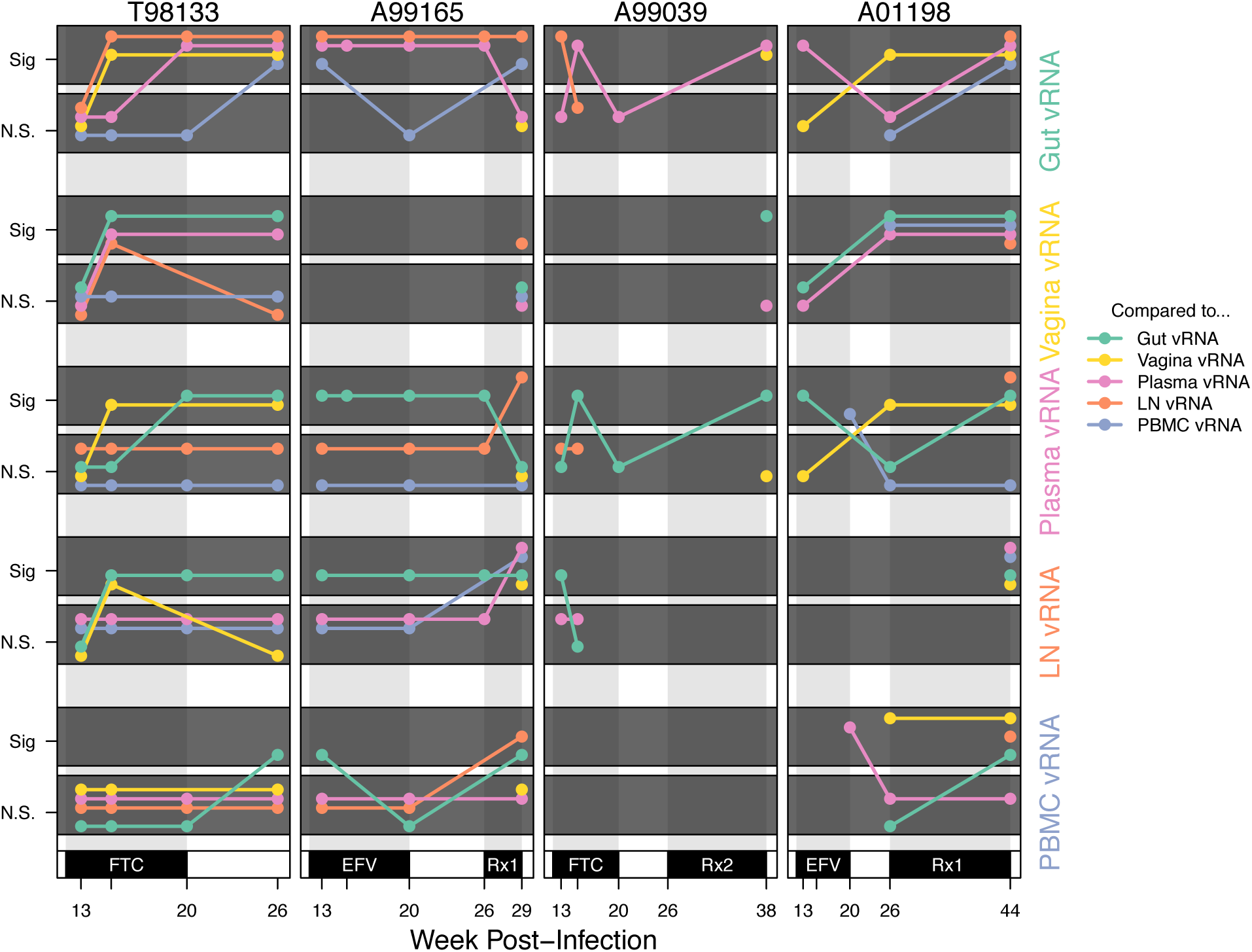
Compartmentalization relationships between different samples reveal transient and stable patterns. Significance results from pairwise compartmentalization tests between vRNA in each compartment for the four macaques (columns) over time (x-axis). Each row represents comparisons between the vRNA in each compartment marked at the right (from top to bottom: gut, vagina, plasma, lymph node, and PBMC vRNA) to all other compartments. Because all pairwise relationships are shown for each compartment, lines are repeated (i.e., the lymph node versus plasma line is present in both the lymph node and plasma rows). K_ST_ tests were significant when they had a p-value below a 5.0*10^-3^ threshold, chosen by setting a 5% false discovery rate, and applying the Benjamini-Hochberg-Yekutieli procedure assuming arbitrary dependence. Coloration indicates that the comparison was done between the focal compartment for the row and lymph node (orange), vagina (yellow), PBMC (blue), gut (green) or plasma (pink). N.S. stands for not significant and Sig stands for significant. Rx1 is treatment FTC+TFV+EFV and Rx2 is the treatment TFV+L870812+DRV/r.

We find that compartmentalization is dynamic through time. While some pairs of compartments in some macaques nearly always appear compartmentalized (e.g., A99165 gut and lymph node vRNA) or never do (e.g., T98133 and A99165, plasma and PBMC vRNA), most often, compartmentalization is transient through time (Figure 5). We also characterized which vRNA samples appeared most similar to each other throughout time (Figure 6). The blood (plasma and PBMC) and lymphoid systems appear well connected with few rejections of being well-mixed in pairwise comparisons. The most compartmentalized comparisons were between the lymph nodes and mucosal tissues (gut and vagina) or between mucosal tissues themselves. Similar results hold if a tree-based test for compartmentalization (the Slatkin-Maddison test) is used instead of a diversity based test (Figure S4 and S5), although the Slatkin-Maddison test is much more affected by sample sizes (Figure S6 and see (24)). Biologically, these observations are in line with expectations about system connectivity.

**Figure 6:**
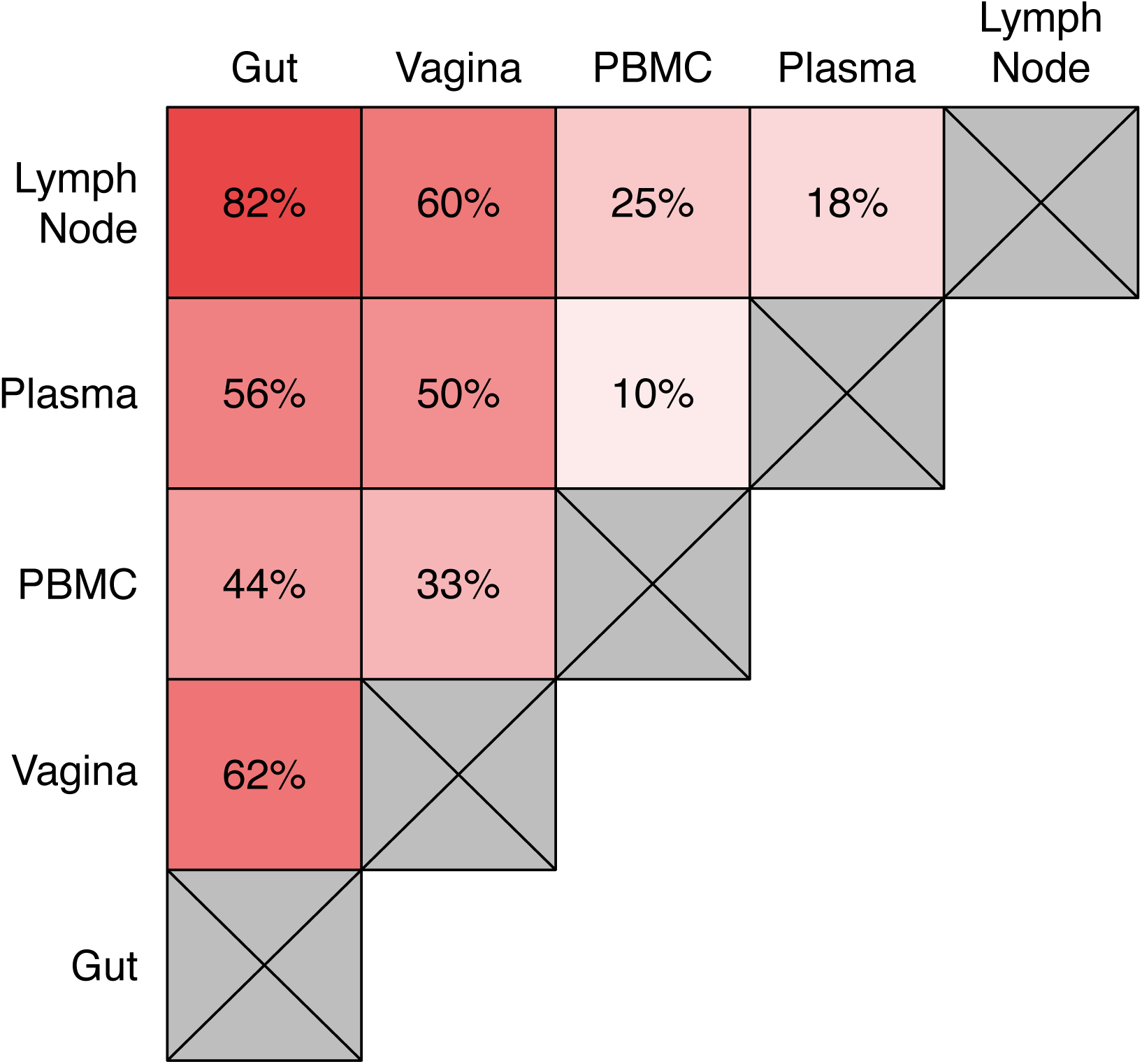
K_ST_ results reveal pairwise RT-SHIV RNA compartmentalization relationships between blood, lymph node, gut, and vagina. Each box lists the percentage of K_ST_ tests between the two compartments that reject the null hypothesis of being well-mixed across all macaques and all time points. Each compartment must have at least 3 sequences from the same macaque and time point for the comparison to be included in the figure.

We asked next if there is a broader relationship between when samples are taken and how compartmentalized they appear. We find that the average pairwise *K_ST_* increases after the onset of drug pressure in all three macaques (Figure 7). This suggests that drug treatment may drive RT-SHIV to become more compartmentalized. K_ST_ then decreases after drug resistance has spread, which suggests that differences between compartments that accumulate during a strong selective pressure may dissipate due to migration once that pressure is removed (i.e., once a DRM has fixed in a population).

**Figure 7:**
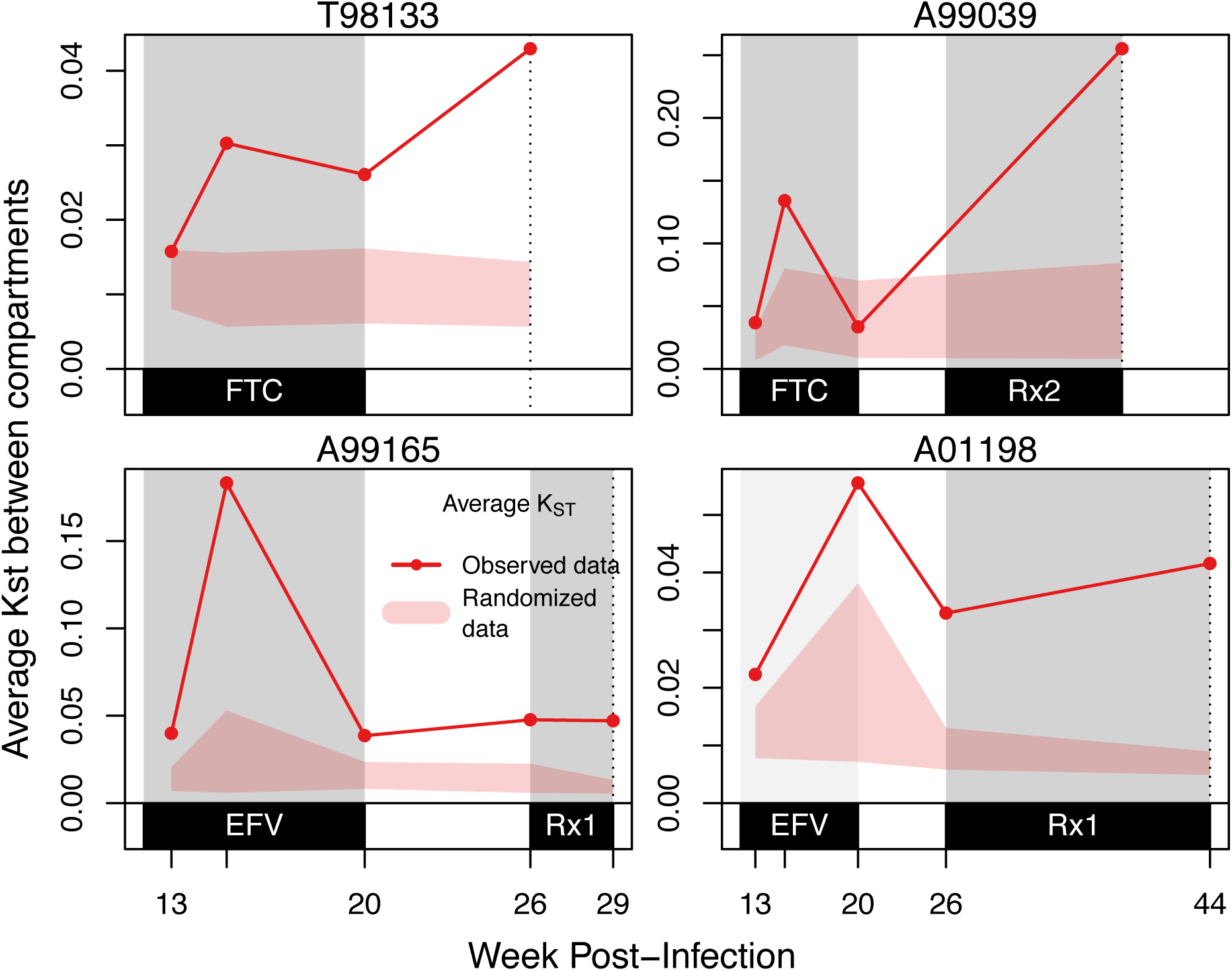
Average K_ST_ increases after the onset of FTC or EFV monotherapy and later declines. For each macaque at each time point, K_ST_ is measured between vRNA in each compartment and the remaining vRNA samples at the same time point (i.e., KST between plasma vRNA versus combined lymph node, vaginal, PBMC and gut vRNA) and then averaged over all compartments present at that time point and plotted (red line). Compartment assignments for sequences are randomized 10^4^ times within each time point (maintaining compartment sample sizes), and the central 95% of randomized K_ST_ values are plotted (pink shading). Grey shading indicates monotherapy or combination therapy, as indicated below the x-axis. Black dotted line indicates the final time point for each animal. Rx1 is treatment FTC+TFV+EFV and Rx2 is the treatment TFV+L870812+DRV/r.

### 2.4 The differences and similarities between vRNA and vDNA

We have characterized the dynamics of the vRNA in different compartments, but this may or may not be representative of the viral reservoirs. We therefore also sampled vDNA in all compartments, which allows us to compare vDNA and vRNA from the same location.

There were marked differences in genetic composition between vRNA and vDNA. Most notably, the vDNA retained a higher percentage of WT viruses than the vRNA in T98133 and A99165 (Figures 3 and 4). In A99039, very little wild-type vRNA or vDNA was present, even at the initial time point, and the population is dominated by viruses with synonymous variants (L187L, K223K, L205L and combinations thereof). However, there were many fewer drug-resistant viruses in the vDNA samples than in the vRNA samples, suggesting that, like the other monkeys, the vDNA is changing more slowly than the vRNA (Figure S3).

To more directly test whether the rate of change was higher in vRNA than vDNA, we computed K_ST_ between consecutive time points separately from both the vRNA or vDNA from the same compartment of each macaque (Figure 8). All pairs of adjacent time points were treated equivalently, even though they were separated by different amounts of time. However, because similar proportions of long and short time differences were present between vRNA and vDNA, we could still compare the number of significantly compartmentalized comparisons as we did in Figure 6. We find that 24/45 (53%) comparisons among consecutive vRNA time points from the same compartment were significantly compartmentalized (indicating a different viral composition), but only 16/51 vDNA comparisons were significantly different (31%). This confirms that vDNA changes more slowly than vRNA. These comparisons also revealed that vRNA from successive time points in the gut were more often compartmentalized than those from the plasma (10/12 comparisons significant compared to 4/15, p = 0.01).

**Figure 8:**
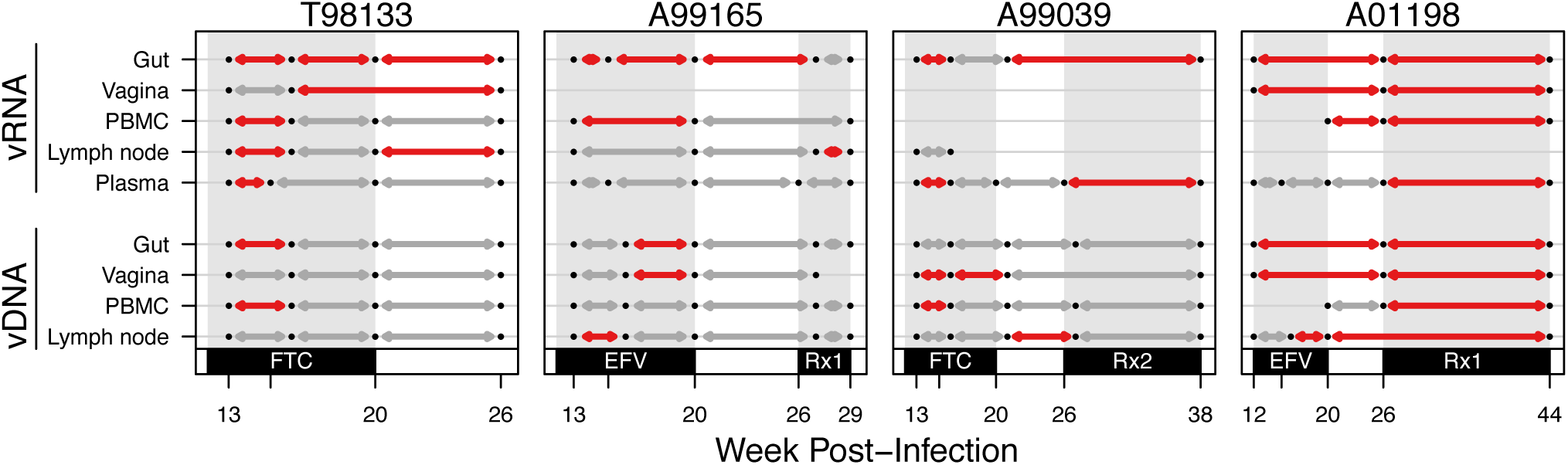
Change in compartmental viral composition over time varies by compartment and drug pressure. For each macaque, K_ST_ tests are plotted between all time points with adjacent samples (i.e., week 12 or 13 compared to week 15 or 16) for each compartment (vRNA top, vDNA bottom). Black dots indicate time points with samples of at least 3 sequences. Red lines indicate that a K_ST_ test comparing those two samples is significantly different (with a 5% FDR). Grey lines indicate a failure to reject the null hypothesis of being well-mixed at the 5% FDR level. Grey shading indicates monotherapy or combination therapy, as indicated below the x-axis. Rx1 is treatment FTC+TFV+EFV and Rx2 is the treatment TFV+L870812+DRV/r.

We then characterized the relationship between the sampled vRNA and vDNA variants by examining viral variants found in the vRNA but not the vDNA and vice versa. We compared all time points for which there were vDNA and vRNA samples *(n* ≥ 3 for both vRNA and vDNA) from a single compartment in a macaque. The proportion of vRNA observed without corresponding vDNA was 25% on average across macaques (Figure S7), suggesting a large portion of the vRNA was transcribed from vDNA at a frequency below the limit of sampling (< ≈1/30). A similar proportion of vDNA is sampled without any concurrent vRNA. This may mean that the provirus is latent (i.e., not currently producing vRNA) or replication incompetent.

In three of the four macaques (T98133, A99165, A99039) vDNA became less representative of the vRNA over time (i.e., a larger proportion of vDNA was sampled without vRNA) (Figure S7), although this effect appeared to plateau in A99165 and A99039. This is consistent with the vDNA building up a "fossil record" or "seed bank" of proviruses that are not transcribed at a given time but that may later be reactivated.

## 3. Discussion

In this paper, we present high-resolution sequencing data from a SIV-HIV chimeric virus (RT-SHIV) sampled across space and time in infected macaques. Although our data is sampled from RT-SHIV-infected macaques, this system corresponds well to HIV-1 in humans in many ways. First, when untreated, RT-SHIV in macaques plateaus at a similar set point as HIV-1 in many humans, suggesting that census population sizes are similar between the two systems. Second, RT-SHIV reaches levels of genetic diversity that are similar to levels in HIV-1, as quantified by average pairwise diversity (16). Third, drug resistance arose via the same DRMs that are common in HIV-1 (M184V/I and K103N) and, fourth, like in HIV-1, the mutations spread via soft sweeps after occurring independently on many backgrounds (23, 25). These observations suggest that an intra-host RT-SHIV population closely mimics that of a intra-patient HIV-1 population undergoing monotherapy, which may occur during drug nonadherence.

Similarities between RT-SHIV in macaques and HIV-1 in humans make RT-SHIV a good model system for HIV-1. In addition, it is possible to sample RT-SHIV much more intensively, including from multiple, longitudinal tissue biopsies, than HIV-1. Thanks to viral RNA and viral DNA single-genome sequences from different time points and different compartments, we have detailed information about evolution in time and space that allows us to make more detailed observations about the dynamics of the system. We have data from 4-5 sampled time points from blood, gut, lymph node, and vagina (both vRNA and vDNA) from four infected macaques. This level of detail allows us to explore several open questions concerning intra-host spatial dynamics and drug resistance.

First, we can ask how well virus from the blood plasma represents the remainder of the intra-host population. This is important because often only the blood is sampled in HIV-infected individuals. We find that while the blood plasma often has the same viral variants as other compartments, it is not necessarily representative of the dynamics occurring in other compartments. For example, the virus population in the gut changes faster than that in the plasma, both among the vRNA and the vDNA. This may be because the CD4+ T cell composition of the gut is different than that of the blood. Specifically, the gut has a higher proportion of effector memory cells CD4+ T cells (T_EM_) than the blood or lymphoid tissues (26, 27). Infection with a CCR5-tropic SIV, including the SIV_mne027_ backbone used in this study (28), leads to significant loss of CD4+ T_EM_ cells in mucosal tissues, such as the gut (29, 30). These CD4+ T_EM_ cells are also shorter-lived than the CD4+ central memory T cells (T_CM_) found at higher frequencies in the plasma and lymphoid tissues (29, 31). This may explain the faster rate of change in the expressed virus (vRNA) and in virally infected cells (vDNA) in the gut. More broadly, this suggests that while surveying the plasma may accurately reveal the types of mutations present in the HIV-1 quasispecies, it does not necessarily reflect the frequency or dynamics of these mutations in other compartments.

Second, we sampled RNA and DNA, which allows us to observe not just the part of the viral population that is actively replicating but also the part of the viral population that is not expressed and may be latent (DNA not represented in RNA). This is of interest because it is not well known how the viral reservoir changes in patients undergoing monotherapy. Previous studies of HIV-1 or RT-SHIV populations during suppressive combination antiretroviral therapy (ART) have shown that the reservoir changes very little, suggestive of little or no ongoing replication during ART (32-35). While studies of actively replicating RT-SHIV populations have suggested that the reservoir is changing through examining changes in vRNA (34, 36, 37), this study directly measures changes in vDNA over time. We find that while new vDNA is produced, the viral composition of the reservoir still remains relatively stable compared to the vRNA. While the majority of vRNA from different compartments of the macaques can contain DRMs, only a small proportion of the vDNA acquires drug resistance, and the majority remains wild-type or genotypes that were present before treatment. This observation suggests that only a small proportion of the vDNA is actively producing virus, and the majority is defective or latent, even during active replication.

Third, our detailed sampling over space and time helps explain results from existing studies on compartmentalization in HIV-1. As summarized in Table 1, with the exception of the central nervous system, most studies that characterize within-patient population structure find that evidence for compartmentalization is found for some comparisons, and not for others. In addition, those studies that sample at different points in time often observe mixed results over time (5-7). Similarly, our data also show that compartmentalization is a transient occurrence in the blood and tissues we sampled, that arises as the population adapts and declines through frequent and fast migration. This transiency may contribute to the equivocal findings across HIV-1 compartmentalization studies. In fact, studies that found either ample support for compartmentalization or no evidence for compartmentalization may have found the opposite result had they sampled at a different time.

Fourth, because our study sampled both across time and from multiple compartments, we were able to determine when and where compartmentalization is most likely. We find that particular compartments, specifically the blood and the lymph nodes, usually harbor the same mutant frequencies, likely because of high rates of migration of infected cells between those compartments. We rarely find evidence for compartmentalization between those compartments. On the other hand, some compartments, specifically the blood and the mucosal tissues, harbor significantly different mutant frequencies, likely because of limited migration of cells between those compartments, such as CD4+ T_EM_ and tissue resident T cells. We usually observe evidence for compartmentalization between those compartments. In addition, we find some evidence that the onset of a selective pressure may cause more compartmentalization, if variants under positive selection within each compartment are increasing in frequency faster than migration can spread them across compartments. This localized adaptive signature might then degrade once selected variants are at high frequency throughout the macaque and migration equilibrates the system (38).

In addition to answering these open questions, our data are well suited to provide insight into population genetic parameters of viral populations. Knowing these parameters is important in order to use existing theory to make predictions about HIV-1 evolution, which could help improve antiretroviral therapies. For example, because most theory assumes that beneficial mutations are rare, it is important to know if this holds true for HIV-1, including for the emergence and dissemination of drug resistance. Next to the rate at which mutations occur in the population, the migration rate and selection strength are important parameters in population genetic models. Together, these parameters control the dynamics of mutation frequencies over space and time.

In the RT-SHIV sequence data, we observe that both migration and selection are powerful drivers of allele frequency changes. Selection causes drug-resistant variants to reach high frequencies in the course of just weeks, and migration causes these same variants to be present throughout the body. However, it is interesting to note that neither force overwhelms the other. If selection was much stronger than migration, beneficial variants should reach high frequencies in their originating compartment before migrating to other compartments, but we observe the same beneficial variants in all compartments nearly simultaneously. On the other hand, if migration was a much stronger force than selection, we would expect to see very similar allele frequencies in all compartments. Instead we observe transient but significant compartmentalization, particularly after the onset of a selective pressure. This suggests that while there are instances where selection or migration may be acting more quickly, neither force is much stronger than the other. Since we can estimate the selection coefficient of certain mutations to be around 0.5 (0.41-0.7), migration must be of the same order of magnitude. Taken together, the balanced forces of strong selection and rapid migration create an incredibly dynamic system.

The number of beneficial mutations that arise per generation in the population is an important driver of how fast a population will evolve. This rate (the population mutation rate or *Θ*) is the product of the number of genomes that can mutate (i.e., the population size) and the beneficial mutation rate. If the population mutation rate is much less than 1 mutation/generation, a long period of time will be required for a beneficial mutation to occur. If the population mutation rate is much larger than 1, mutations will happen often and we will observe multiple beneficial mutations spreading simultaneously in the population (i.e., sweeps will be soft). Since the mutation rate is fixed, observing the number of adaptive mutations can inform our estimates of the population size.

For HIV-1, estimates for the census population size are very large (10^7^ – 10^8^ productively infected cells (39, 40)) Given the mutation rate of HIV-1, this would imply a large enough population mutation rate for beneficial mutations to occur on hundreds of backgrounds, and sweeps should therefore be exceptionally soft. We do not observe this in our data, which implies that the census population size does not fully determine population dynamics. It is therefore useful to estimate an 'effective' population size (N_e_) that better captures the number of replicating viruses over a certain timescale. One common approach to estimate N_e_ is by using the amount of neutral diversity, which led to the first estimates of N_e_ in HIV-1 to be < 10^3^ (41). Indeed, when we use population genetic diversity to estimate population size in our data, we also estimate values of N_e_ on this order of magnitude. However, if N_e_ was this small, the multiple beneficial mutations we observe sweeping simultaneously in our data would be extremely unlikely. This shows that similar to the census population size, the diversity-based N_e_ also does not explain the dynamics of drug resistance emergence. We believe that the diversity-based N_e_ captures a longterm effective population size, which may be very small because it is strongly influenced by bottlenecks and selective sweeps, including the strong population bottleneck of initial infection. It may therefore greatly underestimate the size of the population that actually contributes to resistance evolution.

To estimate an N_e_ that appropriately captures the population dynamics at the time of drug resistance emergence, we use the number of origins of resistance, following Pennings, et al (23). Given that we observe 3-4 different drug-resistant backgrounds per population, we estimate that effective population size is on the order of 10^5^, which is much lower than the census population size but higher than the diversity-based effective population size. The estimate is similar to the estimate of the HIV-1 effective population size (23), which further shows how RT-SHIV and HIV-1 have similar dynamics. Note that our estimate is a lower bound, given that we can not distinguish when the same DRM occurs on the same background repeatedly, and so are likely undercounting the number of independent origins of drug resistance. In macaques like T98133, where there is little pre-existing variation before drug treatment, many origins may be missed. In macaques with much pre-existing variation, such as A99039, our estimates for the number of independent origins are likely to be more accurate, given that each drug-resistant genotype contains several unique non-DRMs. It is also possible that we are missing DRMs that land on genomes already possessing very deleterious mutations, and so are unable to spread.

Our findings of soft sweeps also support previous theoretical results about populations undergoing evolutionary rescue (i.e., populations decreasing in size that can only be 'rescued' by a beneficial mutation which make the growth rate positive). Wilson et al. predict that when there is a high probability that the population acquires an adaptive mutation before it becomes very small (i.e., rescue is likely), the adaptive mutation is likely to spread via a soft sweep (42). In the RT-SHIV populations during drug pressure, all three macaques experienced rescue by development of DRMs and all three experienced soft sweeps.

As described above, we believe that the characterization of the RT-SHIV dynamics is relevant for HIV-1 populations, and specifically those HIV-1 populations treated with monotherapy or ineffective therapy, but this may not extend to suppressive ART. Treating HIV-1 with multiple drugs may cause viral populations to evolve with many fewer beneficial mutations. This can change the dynamics through which drug resistance spreads within a patient. An example of this was previously demonstrated, in which it was shown that more effective combinations of drugs decrease the number of origins of drug resistance mutations at treatment failure (25). Further studies will be needed to determine if the level of dynamism that we observe here is also found in HIV-infected individuals treated with combinations of effective drugs.

Future studies have much to contribute to our growing understanding of intra-patient evolutionary dynamics. Single-genome sequencing provides full linked genotypes, which allow for the evolutionary origin of new mutations to be tracked. However, next generation sequencing could provide a greater depth of data with more sensitive allele frequency estimates, which in turn could be used to better differentiate population compositions across compartments. In addition, isolating and sequencing different types of CD4+ cells could reveal that different cell types form sub-compartments within those examined in this paper, as previous studies have suggested they might (Table 1). To determine the physical source of DRMs, samples would likely need to be taken on shorter intervals and at greater depth to reveal very low frequency variants. Finally, in order to be able to observe additional origins of drug resistance, it may be possible in the future to use barcoded initial viral strains (43).

Our study reveals a dynamic intra-patient evolutionary process in RT-SHIV, featuring abundant mutations, fast migration, and strong selection. As we strive for better care for HIV-infected individuals, a thorough understanding of this process will be essential. RT-SHIV is a powerful tool for quantifying how and where drug resistance emerges, and locating potential drug sanctuary sites, which may accelerate the evolution of resistance. Further, the ability to sample in space presents new opportunities to characterize the latent reservoir and ultimately develop strategies to develop a cure.

## 4. Materials and Methods

### 4.1 Ethics Statement

All animal-related work was conducted according to the Public Health Services Policy on Humane Care and Use of Laboratory Animals (http://grants.nih.gov/grants/olaw/references/PHSPolicyLabAnimals.pdf). Washington National Primate Research Center (WaNPRC) is accredited by the Association for the Assessment and Accreditation of Laboratory Animal Care (AAALAC) International and registered as a USDA Class R research facility. WaNPRC is certified by the NIH Office of Laboratory Animal Welfare (OLAW A3464-01). All animal-related experiments were performed under protocol 2370-25, approved by the University of Washington Institutional Animal Care and Use Committee. Animals were under the care of a licensed veterinarian and all efforts were made to minimize animal pain and suffering, in accordance with the recommendations of the Weatherall report (http://www.acmedsci.ac.uk/viewFile/publicationDownloads/1165861003.pdf). Peripheral blood was collected by venipuncture when animals were under sedation to relieve pain and suffering. Biological samples were collected and transported according to all relevant national and international guidelines.

### 4.2 Animals

Four female pigtailed macaques (labeled T98133, A99165, A99039, and A01198) were infected intravenously with 1 × 10^5^ infectious units of RT-SHIV_mne0_2_7_, a simian immunodeficiency virus (SIV_mne0_2_7_) encoding HIV-1_HxB_2 RT (14, 15) and hereafter referred to as RT-SHIV. The animals were negative at study initiation for serum antibodies to HIV type 2, SIV, type D retrovirus, and simian T-lymphotropic virus type 1. All procedures were conducted while the animals were sedated with intramuscular injection of Telazol (tiletamine/zolazepam, 3-6mg/kg) for minor surgeries (e.g. biopsies) or with Ketamine (10-15 mg/kg) for blood draws and other minor procedures. Animals that were diagnosed by the veterinarian to be experiencing more than momentary pain and distress were evaluated and treated with appropriate analgesic drugs as indicated. The end point of the study was euthanasia at the protocol-specified time point by intravenous Euthasol (80 mg/kg sodium pentobarbital and 10 mg/kg sodium phenytoin) in the saphenous vein after sedation with 3 mg/kg Telazol. (We'll have SL Hu double-check this info.)

Daily monotherapy was administered for 8 weeks between 12-20 weeks post-infection. T98133 and A99039 received FTC subcutaneously (s.c.; 40 mg/kg) and animals A99165 and A01198 received 400mg EFV orally (p.o.). After a 6 week treatment interruption, daily combination ART was administered until necropsy. T98133 and A99039 received suppressive ART consisting of tenofovir (20 mg/kg s.c.), L-870812 (100mg p.o. twice daily), and darunavir/ritonavir (375 mg/50 mg p.o. twice daily). A99165 and A01198 received recycled therapy consisting of EFV (400 mg p.o.), FTC (40 mg/kg s.c.), and tenofovir (20 mg/kg s.c).

Blood was obtained weekly or bi-monthly for peripheral lymphocyte counts, peripheral blood mononuclear cells (PBMC) pellets, and plasma vRNA quantification, as previously described (44-46). The lower limit for quantification was 30 vRNA copies/ml plasma. Lymph node (LN), rectal (gut), and vaginal biopsies were taken at multiple time points throughout the study. CD4+ T cells were isolated from LNs by magnetic separation (Miltenyi). Plasma, PBMC, LN CD4+ T cells, and tissue biopsies were stored at −80 C.

### 4.3 Viral DNA and RNA Isolation and Single-Genome Sequencing (SGS)

Viral RNA and viral DNA were isolated from frozen samples as previously described (47). SGS was performed on vRNA and vDNA as previously described (15). Drug resistance mutations (DRMs) were determined from the IAS-USA 2015 Update of the Drug Resistance Mutations in HIV-1 (48). DRMs listed for drugs not being used to treat macaques were not assessed as DRMs.

### 4.4 Antiretroviral Drug Concentrations

Drug concentrations were quantified by LC-MS/MS analysis using methods similar to those previously published (49, 50). Tissues were homogenized in 1mL of 70:30 acetonitrile-1mM ammonium phosphate (pH 7.4) with a Precellys 24 tissue homogenizer (Bertin Technologies, Montigny-le-Bretonneux, France). Plasma underwent protein precipitation with an organic solution containing a stable-labeled internal standard. For all compounds, a Shimadzu high-performance liquid chromatography system was used for separation, and an AB SCIEX API 5000 mass spectrometer (AB SCIEX, Foster City, CA, USA) equipped with a turbo spray interface was used as the detector. Samples were analyzed with a set of calibration standards (0.02 to 20 ng/mL for tissue and 5–5,000ng/mL for plasma) and quality control (QC) samples. The precision and accuracy of the calibration standards and QC samples were within the acceptable range of 15%. Tissue density was assumed to be 1.06 g/cm3. All LC-MS/MS data underwent QC by a designated individual not directly involved in this study to ensure accuracy.

### 4.5 Muller Plots

Muller plots show the change in frequency of different sampled genotypes over time separately by compartment and macaque. Time points were included in which at least five sequences of vRNA or vDNA were available in a compartment within a macaque (black vertical lines). Time points with fewer than five sequences of vRNA or vDNA in a single compartment were excluded from the plot. Genotypes were called using all positions with minor allele variants at frequency higher than 1% within each macaque (across all compartments and time points). Genotypes with at least one DRM are marked with black hatching.

### 4.6 Assessing Viral Compartmentalization

Compartmentalization of viral genotypes and change in their frequencies over time is assessed using *K_ST_* (51). *K_ST_* is computed as follows:

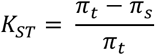

where the total average pairwise diversity of the sample is given by

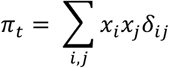

and the average pairwise diversity within each of *k* compartments is given by

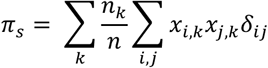

and *x_i,k_* is the frequency of genotype *i* in compartment *k, X_i_* = ∑*_k_ x_i,k_* is the frequency of genotype *i* across all compartments and *δ_ij_* is the percentage of sites at which *i* ≠ *j*.

The significance of *K_ST_* was assessed via permutation test, in which compartmental identities were repeatedly reassigned without replacement, but compartment sizes were maintained. Labels were permuted 10,000 times, and significance was assessed with a one-sided test. The Benjamini-Hochberg-Yekutieli procedure (assuming arbitrary dependence) was used to define a multiple testing significance cutoff that maintained a 5% false discovery rate (FDR).

K_ST_ was used to assess compartmentalization both between two different compartments at the same time point, and between subsequent time points taken from the same compartment (following Achaz, et al (52)).

For verification, compartmentalization tests were also performed using the Slatkin-Maddison test (53) with 10,000 permutations as implemented in HyPhy (54) from trees constructed via UPGMA.

### 4.7 Estimating selection coefficients for DRMs

Selection coefficients were estimated by assuming logistic growth of the drug resistant variants starting at the first treatment administration time point (week 12). This relies on the assumption that DRMs had reached establishment frequency (1/Ns) before treatment. We assumed generation times of 1/day, similar to HIV-1 and previously published (17) and effective population size *(N_e_)* of 1.5*10^5^ (23) and then logistic growth of the allele frequency *x* as follows:

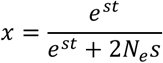

where *t* is measured in generations. We fit s values using two time points – absence (or near absence) of DRMs at week 13 and the DRM frequency at the next available time point (week 15 for T98133 and A99165 and week 16 for A99039). In cases where multiple DRMs were present (e.g., M184V and M184I in T98133), a single s value was fit for both DRMs.

### 4.8 Viral diversity measurements

Intra-compartment diversity was measured for each compartment using the average pairwise diversity (see the definition for π_t_ in **Assessing Viral Compartmentalization** above). The relationship between diversity and time within each macaque was fit with a linear regression of intra-compartment diversity predicted by time (i.e., weeks after infection), separately for vDNA and vRNA. The difference between vRNA and vDNA was assessed by fitting two nested linear models of time on diversity – one in which an indicator variable discriminated between whether a diversity observation was from the vDNA or the vRNA, and one without an indicator variable. An ANOVA was performed to assess the significance of the indicator variable.

### 4.9 Quantifying change in compartmentalization over time

To determine if compartmentalization changed over time, we computed the average *K_ST_* for a macaque at a specific time point as follows: for each compartment at a given sampled week (±1 week), *K_ST_* was computed between the sequences from that compartment and the sequences from all other compartments at that week. This procedure was repeated for each compartment and the resulting *K_ST_* values were averaged to get a measure of total compartmentalization at a given week. This was done separately for each macaque. A permutation test was performed by repeatedly randomizing the sequences at a given time point with new compartmental identities (although maintaining the true compartment sizes) and re-computing the average *K_ST_* statistic 10,000 times.

### 4.10 Estimating effective population size (N_e_) of RT-SHIV

We used both nucleotide level diversity and haplotype structure to estimate the effective population size of the intra-macaque RT-SHIV population. Pairwise nucleotide diversity relates to effective population size N_e_ via the formula:

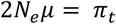

as described in **Assessing Viral Compartmentalization** above. Following Brown (41), we estimated N_e_ using vRNA sequences taken from the blood plasma. It is well known that the estimate of N_e_ is much lower than the census population size, not only in HIV-1 but also in humans and fruit flies.

Aware of the problems with using neutral diversity to estimate N_e_, Pennings et al. identified an alternative approach that should lead to an N_e_ estimate that is much closer to the short term N_e_ that is relevant for drug resistance evolution that can be estimated directly from the nucleotide level encodings of DRMs (23). Briefly, the diversity of different DRM variants *i* through *j* can be summarized by the complement of their summed squared frequencies (or heterozygosity):

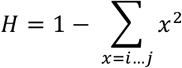

N_e_ can then be estimated from the heterozygosity:

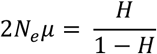

We calculate the heterozygosity using two different definitions of DRM variants: 1) DRM variants that are different by state (e.g., AAU versus AAC for K103N) and 2) DRM variants that can be distinguished by linkage to a common segregating variant (non-DRM mutation at a frequency of at least 10%). We label the resulting estimates of the effective population size N_e,DRM_ and N_e,hap_ respectively. Both N_e,DRM_ and N_e,hap_ are computed among the vRNA variants only and among all sampled variants (vRNA and vDNA) at the first time the vRNA is over 90% drug-resistant, or maximally drug-resistant if 90% drug resistant is never achieved. These times for T98133, A99165 and A99039 are weeks 15/16, week 15 and weeks 20/21, respectively.

All estimates rely on the HIV-1 mutation rate of μ = 1.4*10^-5^ for transitions (such as M184V) and 2 * 10^-6^ for transversions (both forms of K103N) (55).

## 5. Acknowledgements

This work was supported by NIH grant R01 AI080290 (ZA), P51 OD010425 (SLH), P30 AI50410 (ADMK), R35 grant 1R35GM118165 (DAP), and in part with Federal funds from the National Cancer Institute, National Institutes of Health, under Contract No. HHSN261200800001E.

The authors would like to thank the Research Support Group and Virology Core personnel at the WaNPRC for technical assistance and assay support.

The content of this publication does not necessarily reflect the views or policies of the Department of Health and Human Services, nor does mention of trade names, commercial products, or organizations imply endorsement by the U.S. Government.

## Author contributions

Conceived and designed the experiments: AF, SLH, DP, PPe, ZA

Performed the experiments: AF, CK, PPo, BFK

Analyzed the data: AF, MC, ADMK, BFK, SLH, DP, PPe, ZA

Wrote the paper: AF, DP, PPe, ZA

## Supplemental Tables

**Table S1:** Summary of available sequences from RT-SHIV RNA andDNA across compartments separated by macaque and sampling week.

|  | Week | Treatment | Gut |  | Lymph node |  | PBMC |  | Plasma | Vagina |  |
| --- | --- | --- | --- | --- | --- | --- | --- | --- | --- | --- | --- |
|  |  |  | vDNA | vRNA | vDNA | vRNA | vDNA | vRNA | vRNA | vDNA | vRNA |
| T98133 | 13 | FTC | 34 | 28 | 29 | 23 | 33 | 4 | 30 | 21 | 3 |
|  | 15 |  | 29 | 30 | - | - | 29 | 3 | 32 | 24 | 8 |
|  | 16 |  | - | - | 26 | 32 | 31 | - | - | - | - |
|  | 20 |  | 33 | 27 | 24 | 32 | 27 | 11 | 30 | 15 | 1 |
|  | 26 |  | 20 | 32 | 29 | 32 | 29 | 3 | 30 | 26 | 13 |
| A99039 | 13 | FTC | 33 | 13 | 28 | 29 | 26 | - | 26 | 27 | - |
|  | 15 |  | 10 | 4 | - | - | 27 | 1 | - | 16 | - |
|  | 16 |  | - | - | 28 | 8 | 29 | - | 12 | - | - |
|  | 20 |  | - | - | 30 | 1 | 24 | 1 | 27 | 21 | - |
|  | 21 |  | 27 | 7 | - | - | - | - | - | 15 | - |
|  | 26 | TFV+L870812+<br>DRV/r | 11 | - | - | - | 32 | - | 21 | - | - |
|  | 27 |  | - | - | 31 | 1 | - | - | - | - | - |
|  | 38 |  | 29 | 19 | 31 | - | 32 | - | 7 | 19 | - |
| A99165 | 13 | EFV | 29 | 22 | 29 | 29 | 26 | 3 | 29 | 5 | - |
|  | 15 |  | 30 | 27 | - | - | 28 | 1 | 28 | 23 | - |
|  | 16 |  | - | - | 27 | - | - | - | - | - | - |
|  | 20 |  | 28 | 27 | 26 | 23 | 28 | 3 | 21 | 27 | - |
|  | 26 | FTC+TFV+EFV | 30 | 28 | - | - | 30 | 1 | 27 | 4 | - |
|  | 27 |  | - | - | 31 | 32 | - | - | - | - | - |
|  | 29 |  | 35 | 32 | 32 | 28 | 31 | 26 | 27 | 1 | 4 |
| A01198 | 12 | EFV | 19 | 31 | - | - | - | - | - | 22 | 29 |
|  | 13 |  | - | - | 28 | - | - | 1 | 28 | - | - |
|  | 15 |  | - | - | - | - | - | 1 | 3 | - | - |
|  | 16 |  | - | - | 27 | - | - | - | - | - | - |
|  | 20 |  | - | - | 29 | - | 31 | 28 | 30 | - | - |
|  | 26 | FTC+TFV+EFV | 30 | 29 | - | - | 31 | 27 | 31 | 30 | 31 |
|  | 40 |  | - | - | - | - | 31 | - | - | - | - |
|  | 44 |  | 30 | 32 | 29 | 30 | 31 | 31 | 30 | 31 | 30 |

**Table S2:** Linear model fits predicting the change in diversity (average pairwise distance within a compartment (π) over time (week post-infection) separately for vRNA and vDNA by macaque. Model coefficients and their corresponding p-values from the linear fits are given in the table. Significant values at a 5% cutoff are marked with asterisks. The last column gives the p-value from an ANOVA comparing nested models in which vRNA and vDNA by week interactions are fit separately.

|  | vRNA |  |  |  | vDNA |  |  |  | vRNA v.<br>vDNA |
| --- | --- | --- | --- | --- | --- | --- | --- | --- | --- |
|  | (Intercept) | p-value | Weeks | p-value | (Intercept) | p-value | Weeks | p-value | Different? |
| **T98133** | 0.00163 | 0.26112 | 0.00003 | 0.67465 | 0.00093 | 0.56604 | 0.00009 | 0.27531 | 0.6117 |
| **A99165** | 0.00171 | 0.28777 | 0.00008 | 0.34327 | 0.00180 | 0.03372<br>* | 0.00009 | 0.03420<br>* | 0.7161 |
| **A99039** | 0.00505 | 0.00214<br>* | -0.00009 | 0.14265 | 0.00510 | 0.00566<br>* | -0.00005 | 0.47411 | 0.9509 |
| **A01198** | 0.00068 | 0.26715 | 0.00009 | 0.00031<br>* | 0.00286 | 0.00105<br>* | 0.00004 | 0.06563 | 0.0279<br>* |

## Supplemental Figures

**Figure S1:**
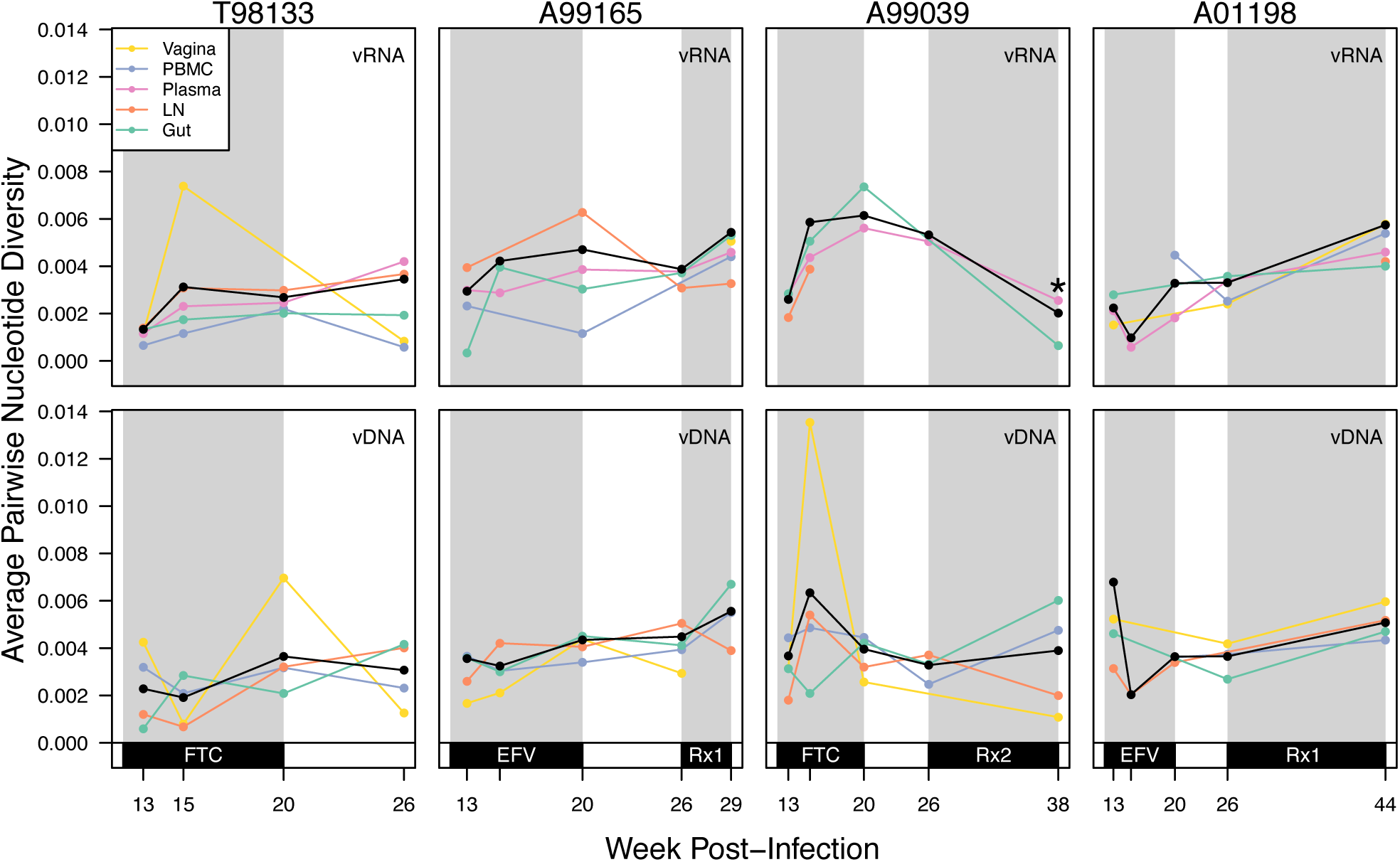
Average pairwise nucleotide diversity within each compartment over time. The average pairwise diversity (π) increases over time in both the vRNA (top row) and vDNA (bottom row) in each macaque (shown in the columns). Each colored line marks a different compartment with the diversity of all compartments combined shown in black. The asterisk marks where diversity decreases at the final sampling point for A99039, and corresponds to viral suppression (see Figure 1). Grey shading indicates monotherapy or combination therapy, as indicated below the x-axis. Rx1 is treatment FTC+TFV+EFV and Rx2 is the treatment TFV+L870812+DRV/r.

**Figure S2:**
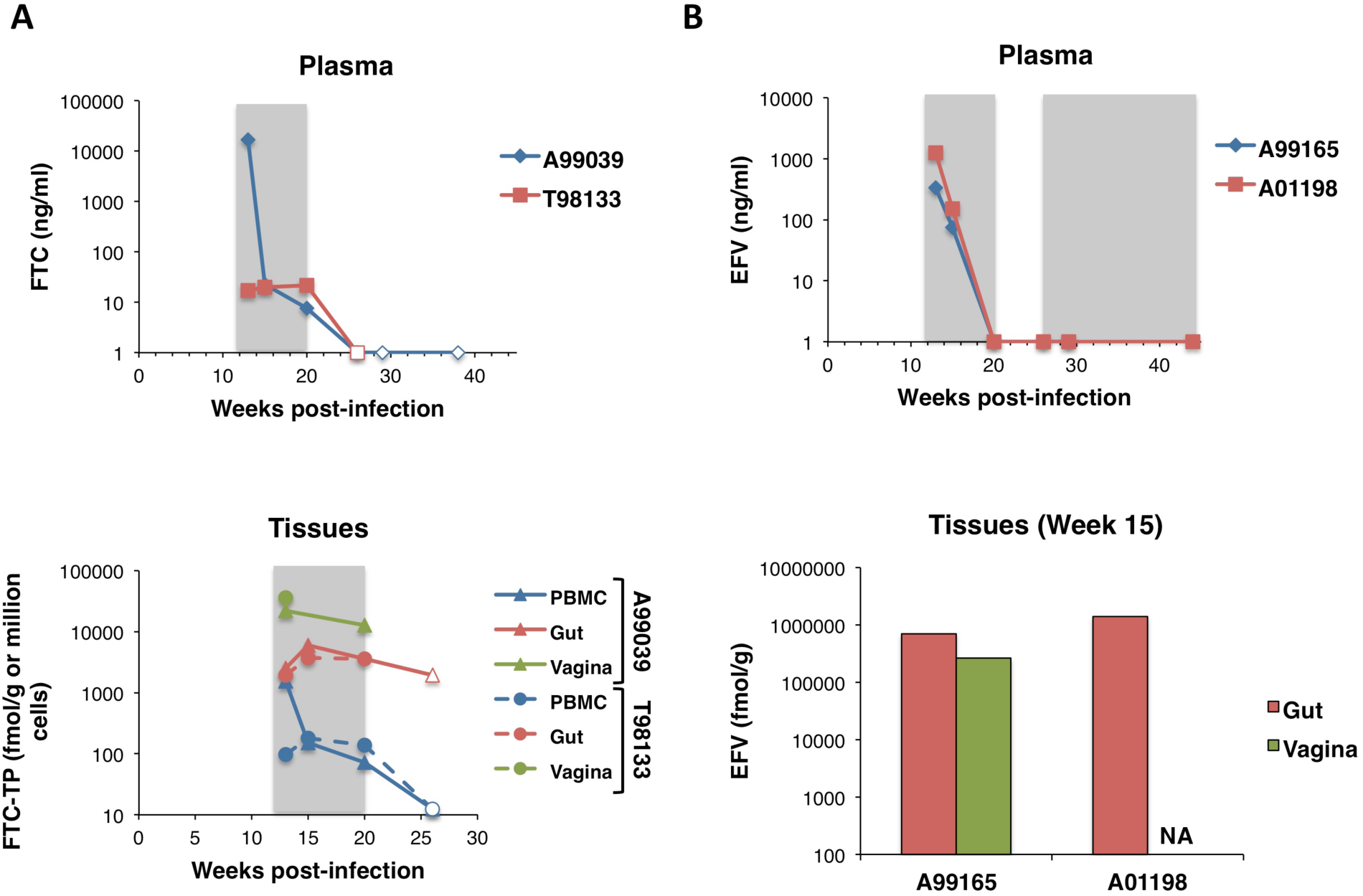
FTC and EFV plasma and tissue concentrations detected during monotherapy. FTC and FTC-TP (A) and EFV (B) concentrations were measured in limited plasma (top) and PBMC and mucosal biopsy samples (bottom) taken from all animals at multiple time points. Grey shading indicates monotherapy (weeks 12-20). Open symbols represent measurements that were below the lower level of quantitation. NA indicates sample was not available. Unfortunately LN tissues were not available for drug measurements.

**Figure S3:**
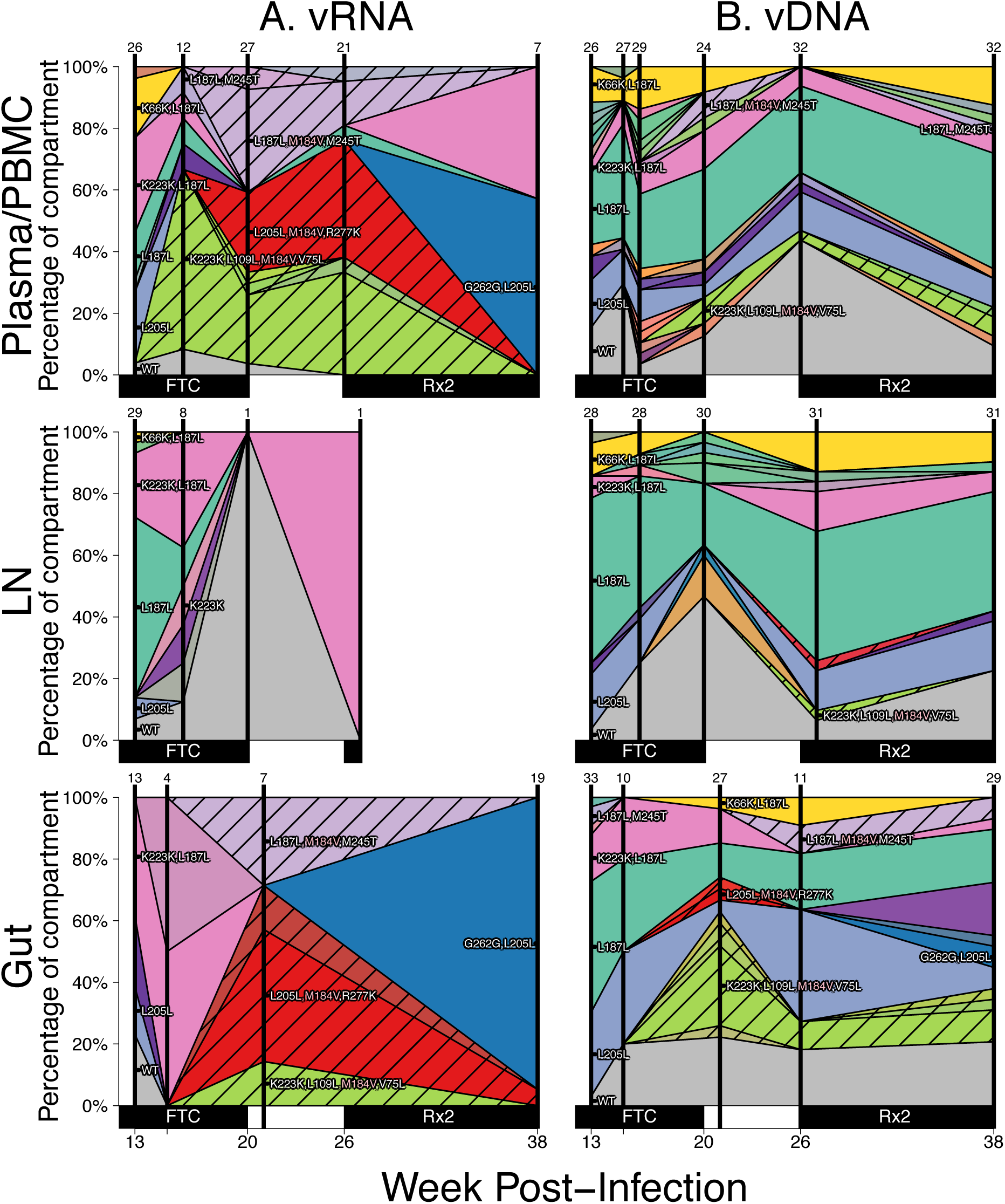
Drug resistance spreads dynamically across several compartments in macaque A99039 over the course of treatment. Figure caption the same as Figure 3. Note, except for WT, the coloration is not preserved between figures (i.e., the yellow in Figure 3 does not represent the same genotype as the yellow in Figure S3). Rx2 is the treatment TFV+L870812+DRV/r.

**Figure S4:**
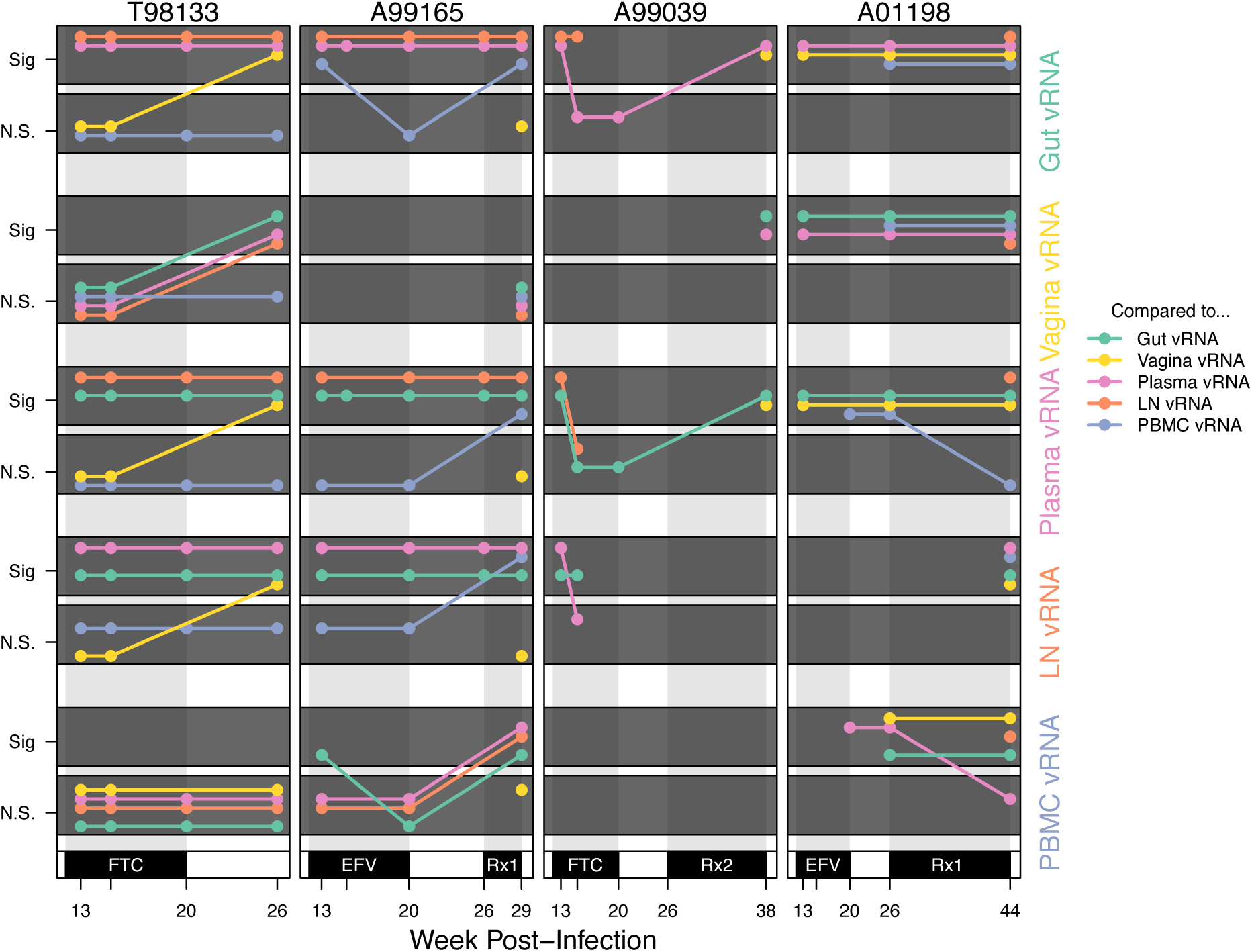
Compartmentalization relationships between different samples reveal transient and stable patterns by Slatkin-Maddison tests. Significance results from pairwise compartmentalization tests between vRNA in each compartment for the four macaques (columns) over time (x-axis). Each row represents comparisons between the vRNA in each compartment marked at the right to all other compartments. Because all pairwise relationships are shown for each compartment, lines are repeated (i.e., the lymph node versus plasma line is present in both the lymph node and plasma rows). Slatkin-Maddison tests were significant when they had a p-value below a 5.0*10^-3^ threshold, chosen by setting a 5% false discovery rate, and applying the Benjamini-Hochberg-Yekutieli procedure assuming arbitrary dependence. Coloration indicates that the comparison was done between the focal compartment for the row and lymph node (orange), vagina (yellow), PBMC (blue), gut (green) or plasma (pink). N.S. stands for not significant and Sig stands for significant.

**Figure S5:**
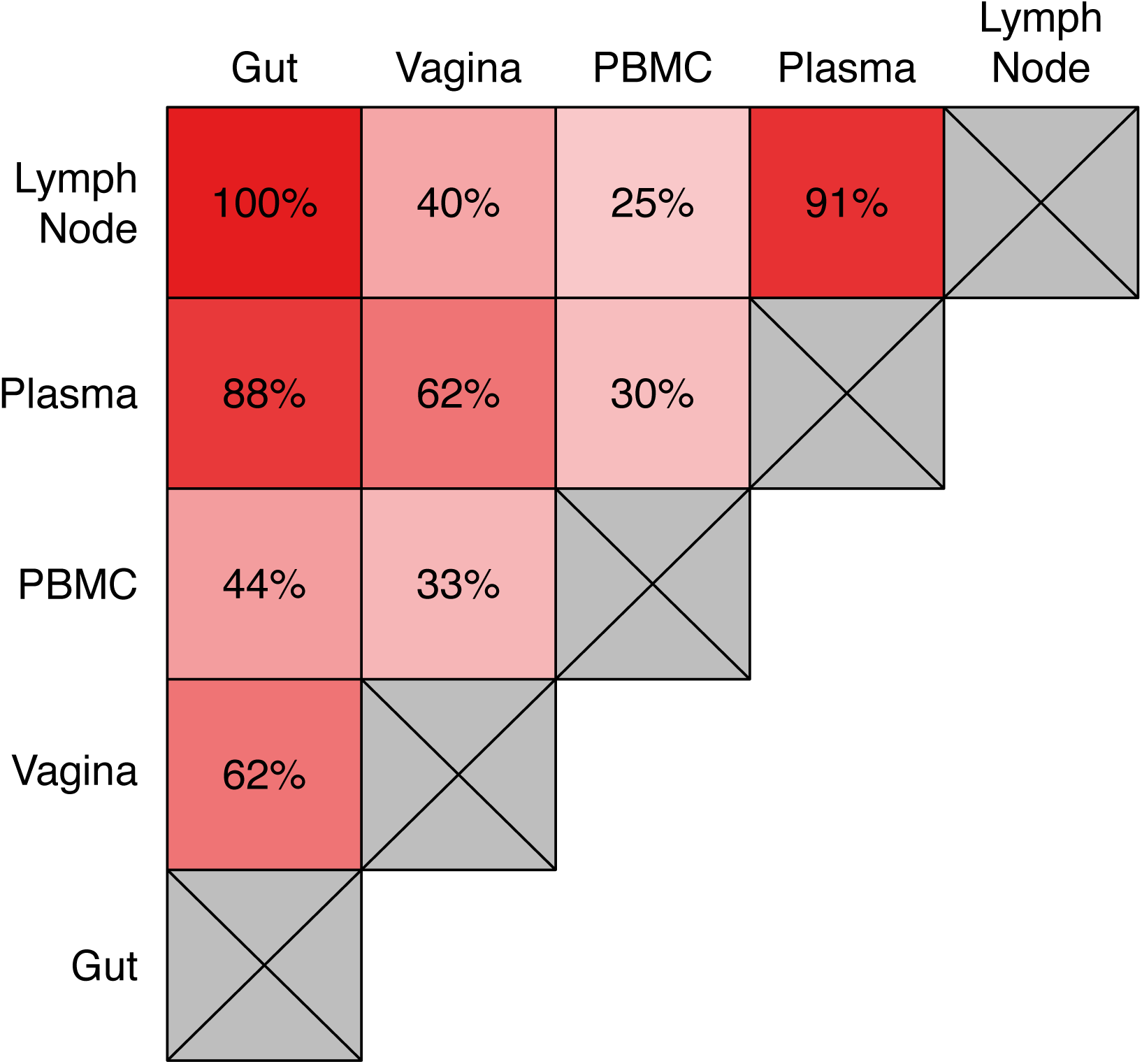
Slatkin-Maddison test results reveal pairwise compartmentalization relationships between blood, lymphoid and mucosal tissue. Each box lists the percentage of Slatkin-Maddison tests between the two compartments that reject the null hypothesis of panmixia across all macaques and all time points. Each compartment must have at least 3 sequences from the same macaque and time point for the comparison to be included in the figure.

**Figure S6:**
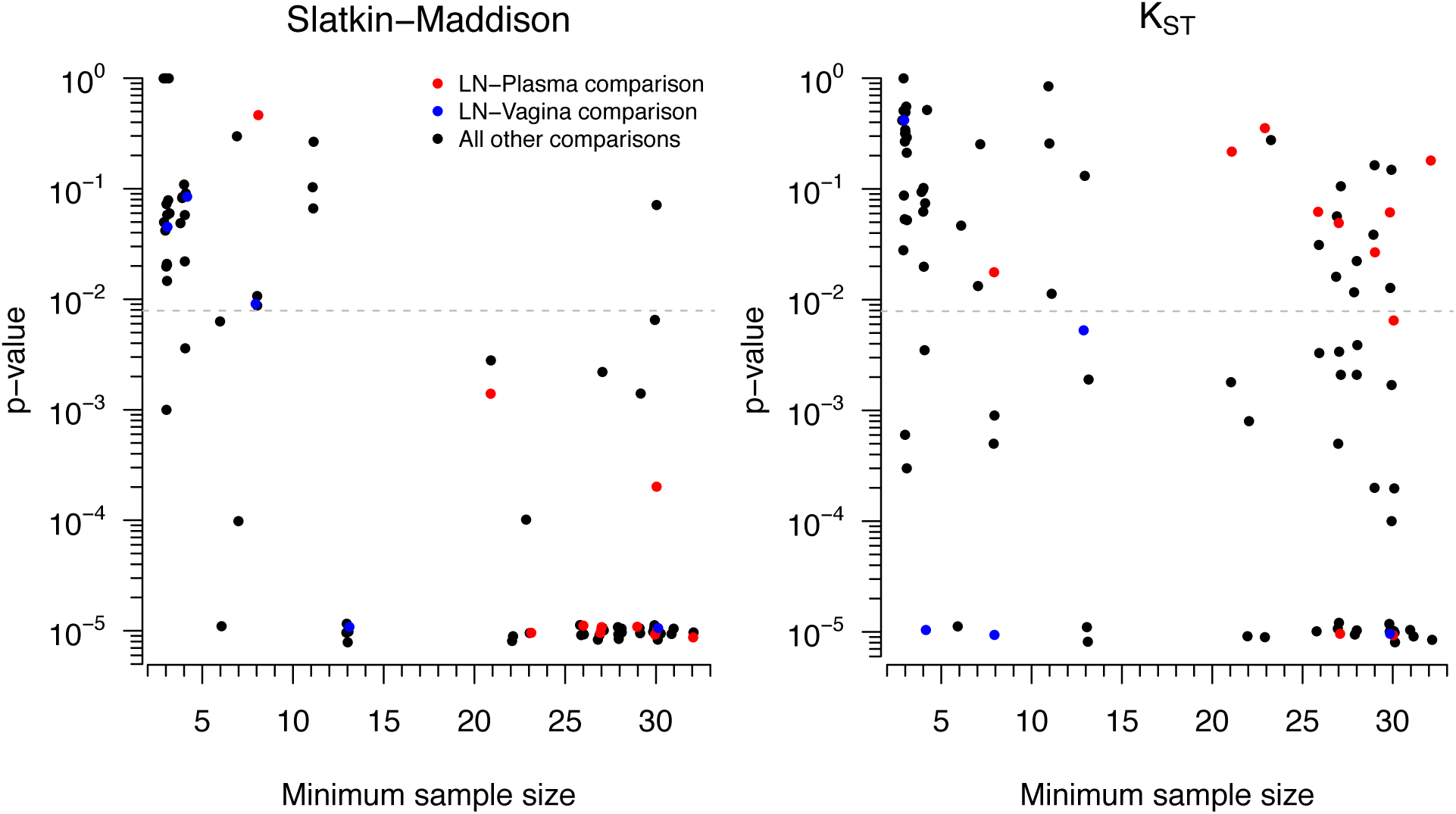
Comparison of Slatkin-Maddison and K_ST_ test p-values and sample size among pairwise vRNA comparisons. For all intra-macaque pairwise vRNA compartmental comparisons at the same time point, the relationship between the minimum sample size (i.e., the smaller of the number of sequences in the compartments of a pairwise comparison) and the p-value for the test statistic is shown using the Slatkin-Maddison test (left) and the K_ST_ test (right). The Slatkin-Maddison test p-values have a greater dependence on sample size (as suggested in Zarate et al (24)) which results in the test having high power when compartment sizes are large (i.e., among LN-Plasma comparisons, shown in red) and having low power when compartment sizes are small (i.e., among the LN-vagina comparisons, shown in blue). A slight jitter is applied to improve point visibility, but each comparison is pictured on both plots.

**Figure S7:**
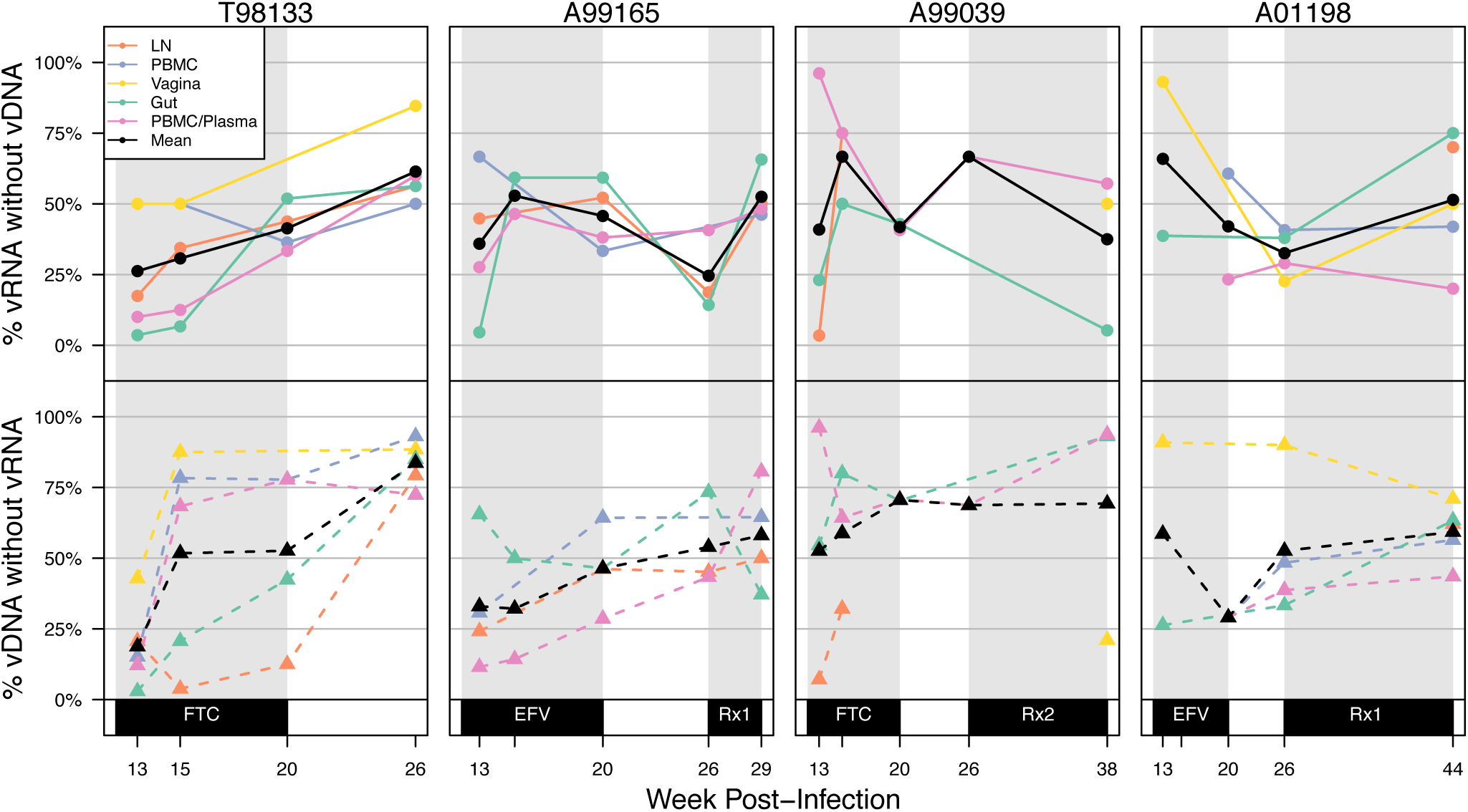
vRNA and vDNA samples from the same compartment harbor variants private to each other. For a given compartment, the percentage of vRNA genotypes without corresponding vDNA genotypes (top) as well as the the percentage of vDNA genotypes without corresponding vRNA genotypes (bottom) is plotted at each time point, in which a genotype was present in both vRNA and vDNA for a given compartment, macaque and time point at a frequency greater than 3 sequences. Plasma vRNA is compared to PBMC vDNA (pink), and all other samples are compared within their sampled compartment (lymph node in orange, vagina in yellow, gut in green and PBMC vRNA verus vDNA in blue). The time point means for both vDNA without vRNA and vRNA without vDNA are shown in black and show overall trends across compartments. Time points that were within a week of each other were considered close enough to compare. Grey shading indicates monotherapy or combination therapy, as indicated below the x-axis. Rx1 is treatment FTC+TFV+EFV and Rx2 is the treatment TFV+L870812+DRV/r.

